# DOMINO: graph diffusion learning identifies spatial domain structures with enhanced accuracy and scalability

**DOI:** 10.64898/2025.12.15.694536

**Authors:** Pan Jia, Nora W Liu, Zixu Ran, Suzanne Maiolo, Tianjun Zhang, Monika Mohenska, Xudong Guo, Cong Wang, Elly Walter, Carmela Ricciardelli, Noor A Lokman, Rebecca Morrow, Martin K Oehler, Jose M Polo, Ning Liu, Fuyi Li

**Author notes:** **Corresponding author** Correspondence to Fuyi Li, Ning Liu, and Jose M Polo.

## Abstract

Spatial transcriptomics enables in situ molecular profiling, allowing to measure the cellular transcriptional output within the tissue. As the tissue architecture is conserved, spatial domains with specific transcriptional patterns can be identified, facilitating the discovery and understanding of functional tissue compartments. Thus, several methods to uncover and identify these spatial domains have been developed. However, most of these existing methods do not scale to rapidly increasing data sizes and focus only on local structure while missing the global view of the tissue. Here, we present DOMINO, a diffusion-optimised contrastive learning framework for spatial domain detection. DOMINO utilises graph diffusion convolution to propagate information beyond immediate neighbours and jointly optimises local and, importantly, global graph structure via contrastive learning. This novel framework yields biologically interpretable domains with clearer boundaries and scales to large datasets, outperforming state-of-the-art methods across healthy and malignant benchmark datasets. We apply DOMINO to a newly generated spatial transcriptomic dataset of endometriosis-associated ovarian cancers, which could not be processed by existing domain detection methods owing to its size. We uncovered conserved proliferative and non-proliferative tumour states that recurred across these tumours and were independently validated in an external clear cell ovarian cancer spatial transcriptomic dataset. Proliferative domains were characterised by elevated expression of *EIF4A1* and *HSPA8*, increased cell cycle activity, reduced mast cell abundance, and coordinated stromal remodelling, including altered fibroblast states and spatial organisation. In parallel, integrative analysis across tumours revealed subtype-specific multicellular ecosystems associated with either endometrioid or clear cell ovarian carcinomas, together with a tumour-excluded stromal domain that could only be resolved through the integration of spatial and transcriptional information. These findings demonstrate how well DOMINO scales up and that it uncovers biologically meaningful spatial programs spanning tumour intrinsic states, tumour microenvironment interactions, and subtype-specific tissue architecture that are not recovered by conventional expression-based clustering approaches.

## Introduction

Spatial technologies are advancing at a rapid pace, revolutionising the way researchers study cells in complex tissues across diverse biological fields, from molecular and developmental biology to immunology and cancer biology ^1;2^. In cancer research specifically, spatial transcriptomics (ST) has become increasingly important as commercial platforms such as Xenium (10x Genomics), Stereoseq (BGI), and CosMx (NanoString) now enable simultaneous measurement of gene expression and cell position within native tissue architecture from both fresh and archival Formalin Fixed Paraffin Embedded (FFPE) tissues at single-cell resolution ^3–5^. By preserving spatial context, ST allows researchers to study how tumour cells and the surrounding tumour microenvironment (TME) interact, uncovering spatially organised programs and spatial domains that drive tumour progression, immune evasion, and therapeutic resistance^3;6;7^. For example, recent studies across diverse cancer tissues have used ST to reveal spatially resolved cell-type heterogeneity that is significantly linked to clinical outcomes or therapy resistance ^8;9^, highlighting the potential of ST as a powerful tool to interrogate spatially resolved regulatory mechanisms that shape tumour behaviour and treatment responses. Accurate interpretation of ST data requires integrating cell position and gene expression simultaneously, which scales up rapidly, increasing statistical complexity and creating significant computational demands, especially in large high-resolution datasets^10^.

Spatial domains are regions of tissue where cells share coherent expression programs shaped by local microenvironments, lineage relationships, or signalling cues^11^. Unlike conventional clustering approaches, which group cells solely on transcriptional similarity, spatial domains capture biologically meaningful tissue compartments defined by both gene expression and spatial organisation. In tumour tissues, identifying spatial domains enables the characterisation of distinct tumour cell states across different regions of the tissue and how these states interact with their TME, e.g. stromal cells, immune cells, etc, providing deeper insights into intra-tumour heterogeneity and tissue architecture. Detecting these domains is therefore critical to understanding tissue organisation, uncovering functional niches, and reconstructing spatial trajectories of disease progression. Recent studies by Kim *et al*. and Singhal *et al*. ^12;13^ have demonstrated the utility of spatial domain analysis in distinguishing tumour from normal tissues, delineating tumour boundaries, characterising tumour–stroma interactions, and mapping immune infiltration patterns, all of which are key determinants of tumour progression and therapeutic response. Consequently, a growing number of computational methods have been developed to identify spatial domains, incorporating spatial information into clustering to infer domain structure directly from ST data, yet challenges remain in accurately resolving complex tissue architecture in large-scale datasets. Supervised approaches such as BuildNicheAssay and hoodscanR leverage known cell type labels to construct neighbourhood profiles ^14;15^. Unsupervised methods, including Banksy and Utag, combine expression with spatial context by aggregating or concatenating neighbourhood information prior to graph clustering ^12;13^. Graph-based deep learning approaches, such as Stagate and GraphST, compute low-dimensional latent representations using graph autoencoders and contrastive learning to capture both local and global structures, thereby enabling more robust spatial domain identification^16;17^. More recently, DECIPHER introduced a cross-scale contrastive learning framework to disentangle intracellular molecular identity from extracellular spatial context, providing scalable cellular embeddings for heterogeneous spatial omics data ^18^. Collectively, these published deep learning-based spatial domain identification methods (Supplementary Table 1) underscore the growing importance of spatial domain detection in resolving biologically meaningful tissue architecture. However, these methods remain limited in scalability and do not consistently preserve accurate domain boundaries, and as spatial transcriptomics datasets continue to increase in size and resolution, accurately capturing both fine-scale local interactions and broader global tissue organisation remains a major challenge.

These challenges are particularly relevant in highly heterogeneous tumour tissues, where resolving spatially organised cellular states and microenvironmental interactions is essential for understanding disease biology. Among solid tumours, ovarian cancer remains one of the most lethal gynecological malignancies, profoundly affecting women’s health worldwide ^19^. Recent work applying ST to high-grade serous ovarian cancers (HGSOCs) has demonstrated the value of such approaches in uncovering novel tumour–immune interactions and stromal architectures that drive disease progression^6^ These studies highlight how resolving spatial organisation at cellular resolution can reveal TME structure that is difficult to infer from dissociated single-cell or bulk profiling. However, the ovarian cancer field has largely centred on HGSOCs, which account for about 70% of epithelial ovarian cancers (EOCs) ^20^. Clear cell ovarian carcinoma (CCOC) and endometrioid ovarian carcinoma (EnOC), which each represent 6 to 10% of EOC cases ^21;22^, are collectively termed endometriosis-associated ovarian cancers (EAOCs) and are often studied as a single disease entity due to their shared association with endometriosis ^23^. Both subtypes are comparatively understudied despite being pathologically, molecularly, and clinically distinct from HGSOCs ^24–26^. Moreover, while CCOC and EnOC share a common association with endometriosis, the extent to which they exhibit shared or distinct spatial architectures and TME organisation remains poorly understood. Insights gained from HGSOCs, therefore, cannot be assumed to generalise to EAOCs, underscoring the need for dedicated spatial studies of these subtypes. Although these subtypes frequently present at an early stage ^24^, stage-stratified outcomes show no significant survival gain ^27^, likely due to reduced platinum sensitivity ^28^. The limited understanding of spatial organisation and signalling programs within CCOC and EnOC continues to hinder biomarker discovery and the development of tailored therapeutic strategies. Accurately resolving spatial domains in these tumours is therefore critical for defining TME architecture and identifying spatially restricted cell states and signalling programs driving disease progression and treatment response.

To accurately identify spatial domains in high-resolution ST data, we developed DOMINO (Diffusion-Optimised doMain IdeNtifier in spatial Omics), a self-supervised learning method based on a multi-view graph contrastive learning framework that robustly identifies spatial domains in large-scale ST datasets. Here, we show that DOMINO not only identifies biologically meaningful tissue domains with state-of-the-art accuracy but also addresses the computational limitations of existing methods, enabling the characterisation of spatial domains. To demonstrate its potential, we applied it to a newly generated cohort of EAOC, which led to the identification of spatial domains underlying EAOC biology.

## Results

### Overview of the DOMINO’s model architecture

The overall architecture of DOMINO is shown in Figure 1. DOMINO is built on a self-supervised multi-view graph contrastive learning framework to jointly model local and global tissue structure to derive biologically meaningful representations. It is designed to integrate spatial coordinates and gene expression data to robustly identify tissue domains from ST data.

**Figure 1:**
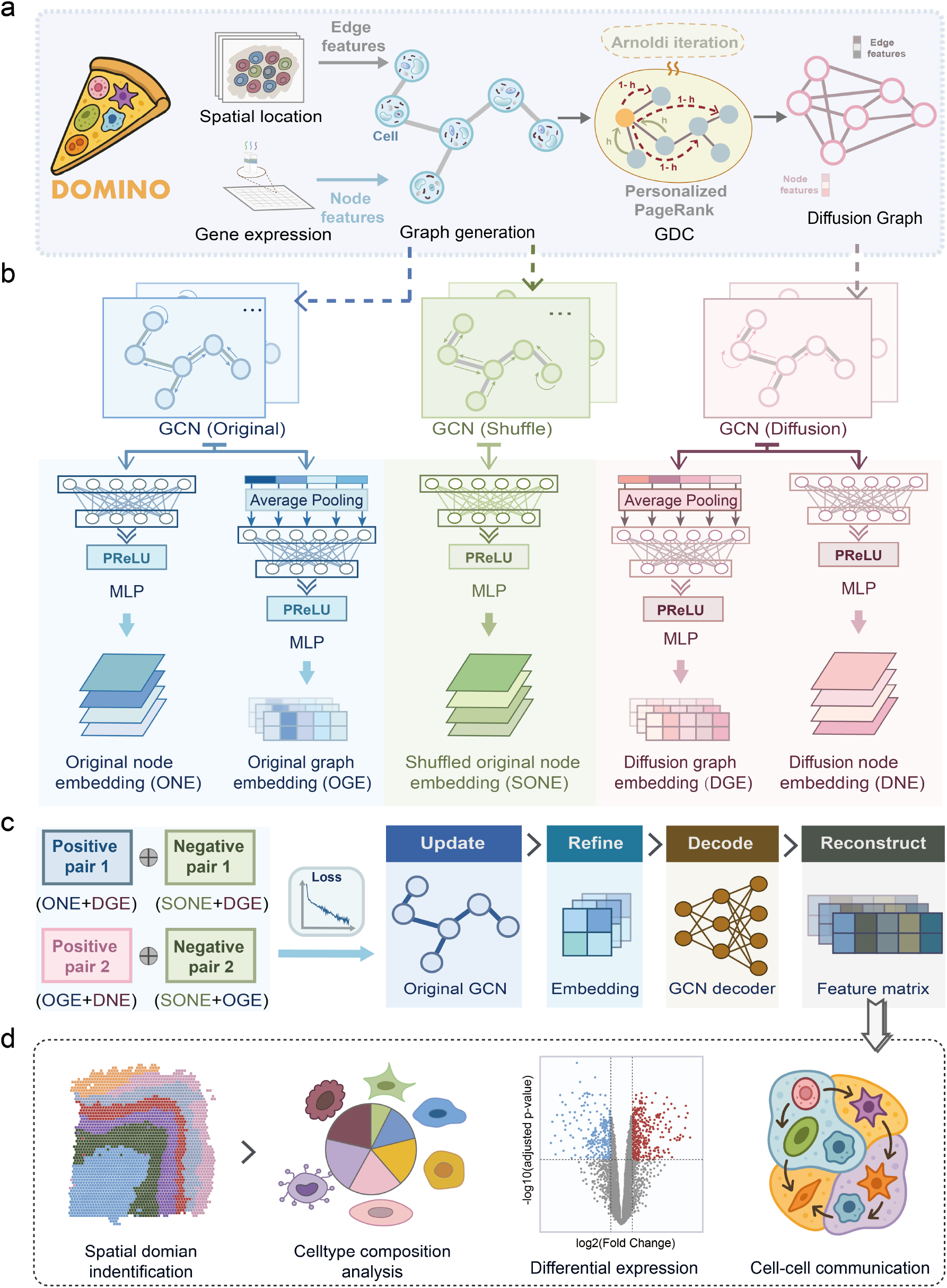
The overall framework of DOMINO. **a.** DOMINO uses spatial coordinates and gene expression of each cell from spatial transcriptomics data as model inputs to construct an undirected graph. A diffusion graph is further derived using graph diffusion convolution with personalised PageRank. **b**. Multiple graph views are encoded through graph convolutional networks. The original graph, shuffled graph, and diffusion graph each generate node-level and graph-level embeddings. These embeddings are used to build multi-view representations in latent space. **c**. A self-supervised contrastive learning framework is applied by defining positive and negative embedding pairs across different graph views and optimising a contrastive loss to refine the encoder. Final embeddings are then decoded back to the original feature space, with a reconstruction loss to ensure that the feature matrix retains spatially informed transcriptomic information. **d**. The learned feature matrix can then be used to identify spatial domains, which can further be used in different downstream analyses, including cell type composition analysis, differential expression, and inference of cell–cell communication.

DOMINO takes both spatial coordinates and gene expression profiles from ST data as input (Fig. 1a). It first constructs an undirected graph where each cell’s gene expression profile is represented as a node, with edges connecting cells based on their local spatial proximity. This graph therefore captures the local spatial architecture by encoding which cells are located next to one another. The main advance of DOMINO is that it then generates an additional graph that propagates information across the whole tissue, so each cell is not only influenced by its immediate neighbours but also by more distant cells. This process is done via diffusion (see Methods). By adding the diffusion graph, DOMINO accounts for both local cell neighbourhoods and the global tissue architecture, enabling accurate and coherent domain detection that mirrors how pathologists look at regions of interest within the context of the whole tissue when performing pathological annotation. Subsequently, both the original and diffusion graphs are passed through graph convolutional network (GCN) encoders (Fig. 1b), which project node features into low-dimensional latent space. In this space, DOMINO then incorporates expression and spatial information using self-supervised graph contrastive learning (Fig. 1b,c) (see Methods). These representations are then decoded to reconstruct the input matrix, enabling calculation of the reconstruction loss.

Finally, reconstructed feature embeddings are then used to identify spatial domains via unsupervised clustering. The resulting labels are spatial domains that form the basis of distinct downstream analyses (Fig. 1d). These include cell-type composition within domains, differential expression analysis between domains and regions, and inference of domain-specific cell–cell communication patterns. By integrating local and global graph information under a self-supervised framework, DOMINO enables accurate and efficient identification of spatial domains.

To assess the contribution of each module in DOMINO’s model architecture, we performed an ablation analysis (Supplementary Fig. 1). Removing the reconstruction module, or using it alone, markedly reduced performance, highlighting the need for joint optimisation with contrastive learning. Removing the diffusion graph or using a single graph view also impaired accuracy, suggesting the benefit of incorporating global context. The complete DOMINO model achieved the highest score in the ablation study, demonstrating the synergistic effect of all its components.

### DOMINO outperforms existing methods with precise boundaries and scalability to large datasets

To evaluate the performance of DOMINO against existing methods, a comprehensive benchmark was conducted across a diverse collection of ST datasets. We first curated datasets generated from five widely adopted ST commercial platforms, including 10X Xenium, NanoString CosMx, BGI Stereo-seq, Vizgen MERFISH, and 10X Visium, which collectively cover the major technological strategies, imaging-based and sequencing-based, in the field. To cover the breadth of tissue contexts, we included both cancer and non-cancer tissues spanning distinct spatial architectures. Cancer datasets have highly heterogeneous TMEs with intermixed malignant, stromal, and immune compartments, whereas non-cancer datasets, such as brain and colon, contain hierarchically organised spatial domains. This combination enabled us to do a rigorous assessment of DOMINO across varying contexts where spatial boundaries are either diffuse and irregular (tumour samples) or anatomically constrained (normal tissues). When selecting datasets, we included datasets frequently used in prior benchmarking studies, allowing direct comparison to established ground-truth labels, including expert pathologist annotations where available, as accurate recapitulation of histopathological structures represents an essential first step in validating spatial domain detection methods. In total, this benchmark experiment encompassed 60 ST datasets (40 full-size and 20 down-sampled) with sizes ranging from 2,695 spots to 555,579 cells (Supplementary Fig. 2 and Supplementary Table 2), providing a comprehensive and balanced resource to evaluate model precision, robustness, and scalability. In particular, because several methods did not scale to all datasets, we also performed benchmarking on down-sampled data where necessary. However, the ultimate goal of spatial domain analysis is not only to reproduce histological annotations but to fully leverage the rich molecular and spatial information captured by ST technologies to uncover finer-scale and transcriptionally informed tissue domains that may not be apparent through conventional histopathology alone.

We focused on benchmarking against methods that reflect the current state-of-the-art in spatial domain identification. Specifically, we included two graph-based deep learning approaches, GraphST and Stagate, both of which have been reported to achieve leading performance in spatial domain recognition ^29;30^. We also included two non–deep learning methods, Banksy and UTAG, which have demonstrated strong performance, particularly in large-scale datasets and in cancer tissues^12;13^.

Compared with ground truth annotations (Fig. 2a–d; Supplementary Figs. 3–14), DOMINO consistently produced spatial domain maps that most closely matched visually, preserving both the overall tissue architecture and fine structural details across diverse platforms. GraphST also achieved strong performance, especially on 10X Visium datasets, but its accuracy declined on single-cell resolution platforms, where domains appeared over-smoothed or fragmented (Supplementary Figs. 3, 6). Banksy performed relatively well on large-scale Stereo-seq datasets, successfully delineating major tissue compartments, although its results were less consistent across other platforms. UTAG demonstrated good scalability but tended to generate noisier or less well-defined boundaries compared to other methods.

**Figure 2:**
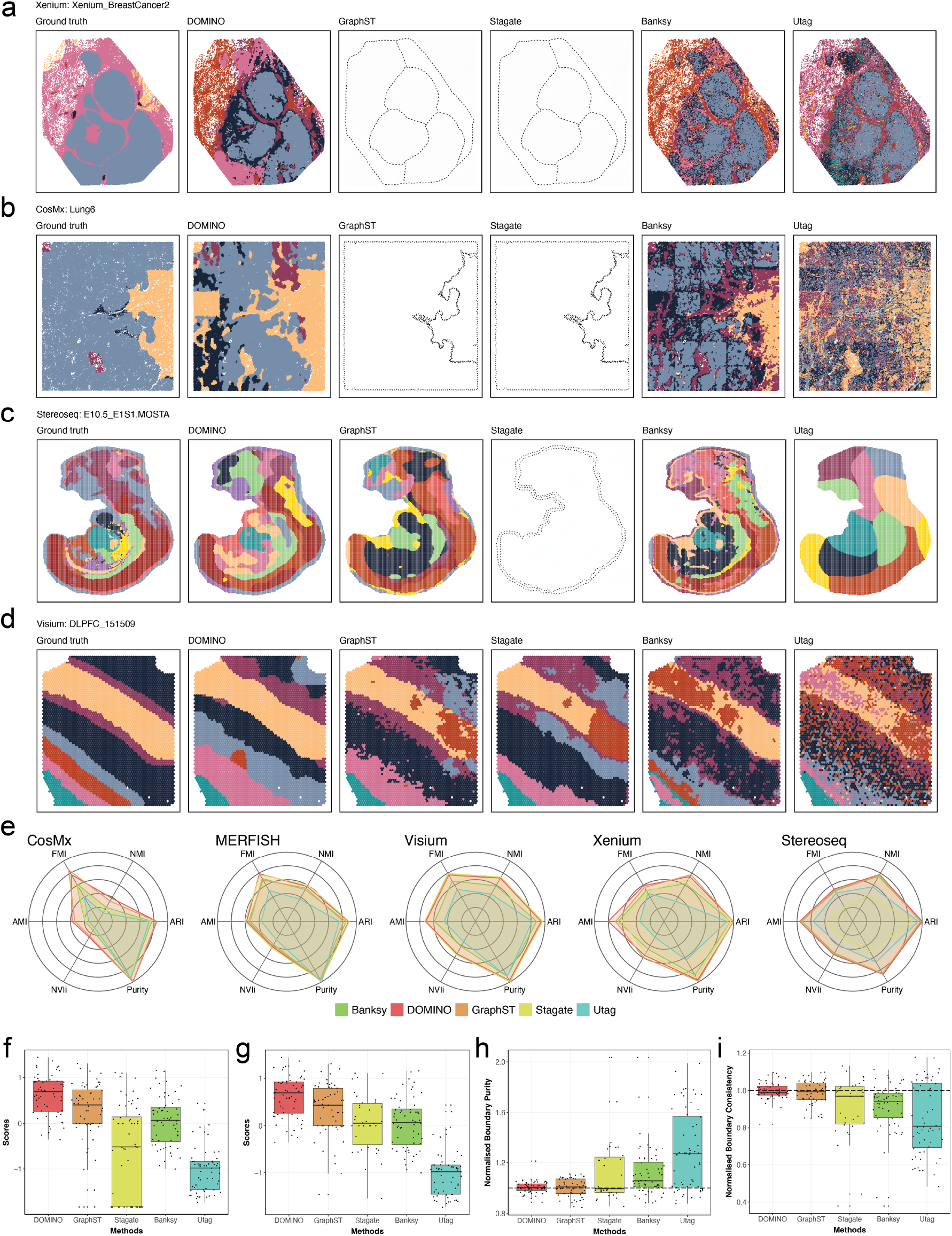
Benchmarking analyses comparing DOMINO against state-of-the-art methods. **a-d.** Visualisation of spatial domain identification across representative datasets from different platforms, with ground-truth annotations shown alongside results from DOMINO, GraphST, Stagate, Banksy, and UTAG. Visualisation with only dotted outlines indicates that the method failed to process the ST data. ST data are **a** Xenium breast cancer, **b** CosMx non-small cell lung cancer, **c** Stereoseq mouse embryo and **d** Visium human dorsolateral prefrontal cortex. **e**. Radar plots summarising six clustering accuracy metrics across five platforms (CosMx, MERFISH, Visium, Xenium, Stereo-seq), including adjusted Rand index (ARI), normalised mutual information (NMI), adjusted mutual information (AMI), Fowlkes–Mallows index (FMI), inverted normalised variation of information (NVIi), and Purity. **f-g**. Summarised benchmarking scores combining multiple accuracy metrics (see Methods) across all datasets. f. penalised methods that failed to process the datasets by assigning the lowest score, while g. Ignore all failure events when calculating the score. **h**. Normalised boundary purity (BP) scores. BP measures local label purity at predicted domain boundaries, with higher raw values indicating fewer mixed labels and sharper boundaries. Scores were normalised to the corresponding ground-truth labels within each dataset; therefore, values closer to 1 indicate boundary purity more similar to the ground truth. **i**. Normalised boundary consistency (BC) scores. BC measures the stability of domain assignments on either side of predicted boundary edges, with higher raw values indicating more coherent regions flanking a boundary. Scores were normalised to the corresponding ground-truth labels within each dataset; therefore, values closer to 1 indicate boundary consistency more similar to the ground truth.

To quantify the performance, we used radar plots to summarise six metrics: Adjusted Rand Index (ARI), Normalised Mutual Information(NMI), Adjusted Mutual Information (AMI), Normalised Variation of Information (NVI), Fowlkes–Mallows Index (FMI) and Purity (see Methods), across different technology platforms, showing DOMINO consistently formed the outermost or co-outermost polygon (Fig. 2e), indicating DOMINO can outperform other methods in accurately identifying tissue domains. The advantage of DOMINO was particularly evident in cancer and single-cell resolution datasets (median ARI of 0.631), where the underlying microenvironment is highly heterogeneous and spatial boundaries are less distinct (Supplementary Fig. 15). In the DPLFC datasets, which are spot-based but commonly used as bench-marking standards, DOMINO (median ARI of 0.733) performed on par with GraphST (median ARI of 0.731) (Supplementary Fig. 16), indicating comparable recovery of large, well-defined tissue compartments in spot-resolution data. Taken together, these analyses demonstrate that DOMINO achieves consistently strong concordance across datasets with either widely accepted or expert-defined domain annotations (Fig. 2f, g), and excels particularly when the tissue structure exhibits sharp or irregular transitions, as often seen in cancer systems.

In addition to clustering-based metrics, we observed that DOMINO delineated spatial domains with markedly clearer inter-domain boundaries compared to other methods (Fig. 2a–d; Supplementary Figs. 3–14). To quantitatively assess boundary sharpness and stability, we developed two complementary graph-based metrics: Boundary Purity (BP) and Boundary Consistency (BC) (see Methods). BP quantifies the average label purity among cells located along predicted domain borders, while BC measures how consistently neighbouring regions on either side of a boundary remain internally homogeneous. Across all datasets, DOMINO produced normalised BP and BC values that were consistently close to the ground-truth reference, with the smallest variation across samples compared to other methods (Fig. 2h, i). This indicates that its predicted domains preserve boundary integrity and consistency comparable to the annotated tissue domains. In contrast, other approaches showed larger deviations from the reference value and greater variability across datasets, suggesting either diffuse, irregular or overly simplified boundaries despite achieving comparable overlap-based scores in some datasets. We further verified that the improved BP and BC values were not primarily driven by post hoc label refinement by evaluating these metrics across DOMINO ablation variants. The full DOMINO model achieved the strongest overall boundary performance, while the refined and unrefined variants showed highly similar BP and BC values (Supplementary Fig. 17), indicating that the observed boundary improvements reflect the learned domain structure rather than artificial boundary smoothing.

To evaluate if DOMINO tends to generate over-smoothed boundaries. We performed boundary- and region-sensitive analyses by estimating boundary-level agreement using boundary F1 and recovery of annotated regions across different spatial scales using best-match recall and IoU (see Methods). DOMINO achieved competitive boundary F1 scores and maintained strong recovery of small and medium-sized regions relative to other methods (Supplementary Fig. 18). These results indicate that DOMINO does not improve spatial coherence by simply merging domains into large blocks, but instead preserves boundary fidelity and fine-scale tissue structures while producing stable spatial domains.

With the rapid growth of ST technologies, data sizes have expanded from thousands to hundreds of thousands of cells per sample, making scalability a critical requirement for domain identification methods in addition to accuracy. For example, we noticed that while DOMINO was able to process all datasets, regarding of the size, Stagate and GraphST exhibited poor scalability, failing to process 29 and 10 full-sized datasets (cell count ranging from approximately 20,000 to 500,000), respectively, and thus yielded incomplete benchmarking results (Fig. 2a–c; Supplementary Figs. 4, 5, 7, 9–11). To systematically assess scalability, we generated simulated ST datasets ranging from 1,000 to 500,000 cells and benchmarked all models under a modern computing environment (14-core CPU, 100 GB RAM, an A800 GPU with 80 GB VRAM). In this experiment, we also included DECIPHER^18^, which is designed for analysing large-scale ST data. As a result, Banksy hit a memory outage with 400,000 cells, while both GraphST and Stagate failed beyond 100,000 cells (Supplementary Fig. 19). While DOMINO, Utag and DECIPHER can successfully process the full range of dataset sizes, DOMINO can outperform the other two in domain identification accuracy (Fig. 2 and Supplementary Fig. 20). Together, these results demonstrate that DOMINO achieves state-of-the-art domain and boundary detection accuracy across diverse datasets while maintaining excellent scalability to large-scale ST data. By overcoming the diffuse-boundary artifacts and computational bottlenecks of existing methods, DOMINO provides a robust and practical framework for domain identification in current and future ST data analysis.

Given that graph-based learning methods may be affected by stochastic training and random sampling, we evaluated the stability of DOMINO by repeating the analysis across five independent random seeds. DOMINO produced consistent metrics across repeated runs, with low variance (0.0005 to 0.001) that was broadly comparable to other graph-based methods such as GraphST and Stagate (Supplementary Fig. 21). This indicates that DOMINO is not unusually sensitive to random initialisation or stochastic sampling, supporting the robustness of the inferred spatial domains under repeated training.

### DOMINO enables spatially informed multi-sample integration

In the benchmarking experiments, we analysed each sample independently. While sample-wise analysis is appropriate for evaluating domain structure within individual tissue sections, multi-sample, especially cohort-level studies require strategies for identifying and comparing corresponding domains across patients or experimental conditions. Because most ST technologies are slide-based, unwanted technical variation, such as batch effects is often inevitable. Therefore, multi-sample domain identification requires methods that not only can scale up significantly, but can also remove unwanted variations while preserving biologically meaningful spatial structure.

Previous studies have addressed this problem by applying non-spatial batch correction methods, such as Harmony ^31^, to integrate multi-sample spatial transcriptomics datasets before or during spatial domain identification^13^. To evaluate whether DOMINO can support joint domain identification across multiple samples, we utilised data from the Visium DLPFC (human brain), MERFISH mouse colon and Stereo-seq mouse embryo datasets. We then applied Harmony to the DOMINO feature embeddings obtained after contrastive learning and then performed joint clustering across samples. We compared this strategy with non-spatial integration, as well as with GraphST, Stagate and Banksy, with and without Harmony-based batch correction using the 6 metrics from the benchmarking experiment. In the DLPFC datasets, where batch effects between samples are minimal, applying DOMINO without Harmony outperforms other strategies by achieving the highest score across all metrics (Supplementary Fig. 22). In datasets with batch effect, joint clustering using Harmony-corrected DOMINO embeddings outperformed others by yielding the strongest performance (Supplementary Figs. 23-24). Taken together, these show that DOMINO supports spatially informed multi-sample integration to remove batch effect while preserving biological variations.

### DOMINO resolves conserved proliferative programs across endometrioid and clear cell ovarian carcinomas

The 10X Xenium 5K panel was used to profile four EAOC tumors, including two endometrioid ovarian carcinomas (EnOC 2193C and 2459B) and two clear cell carcinomas of ovarian (CCOC 2008A and G24D) (Fig. 3a,b, Supplementary Table 3). One endometrioid tumor (2193C) was obtained from a patient with concurrent endometriosis, while the remaining tumors had no clinically documented history of concurrent endometri At a resolution consistent with expert pathological annotation (*i*.*e*., 4 categories), DOMINO consistently separated tumour and non-tumour regions across all cases (Supplementary Fig. 25a). Increasing the resolution to seven domains per sample revealed finer spatial organisation within each tumour, identifying multicellular spatial domains composed of distinct combinations of malignant and non-malignant cell populations. (Fig. 3c, Supplementary Fig. 25b). Consistent with our benchmarking results, GraphST did not scale to the high cell counts of these Xenium datasets and therefore could not be applied to any full tumours. Although Banksy could be run on the complete datasets and broadly separated tumour from non-tumour regions, it failed to further resolve tumour-enriched regions into smaller biologically coherent domains, even when the resolution was increased to generate 20 Banksy subdomains (Supplementary Fig. 25c). To assess whether DOMINO domains reflected genuine biological distinction, we performed Uniform Manifold Approximation and Projection (UMAP) of epithelial cells using gene expression alone. Cells from different DOMINO domains showed partial transcriptional separation on UMAPs across samples, indicating that the identified domains are supported by underlying transcriptional differences despite not being fully distinguishable without spatial context (Supplementary Fig. 25d). These findings suggest that DOMINO captures biologically meaningful epithelial states rather than artefacts of over-clustering and highlight the value of integrating spatial information to resolve intratumoural organisation at cellular resolution.

**Figure 3:**
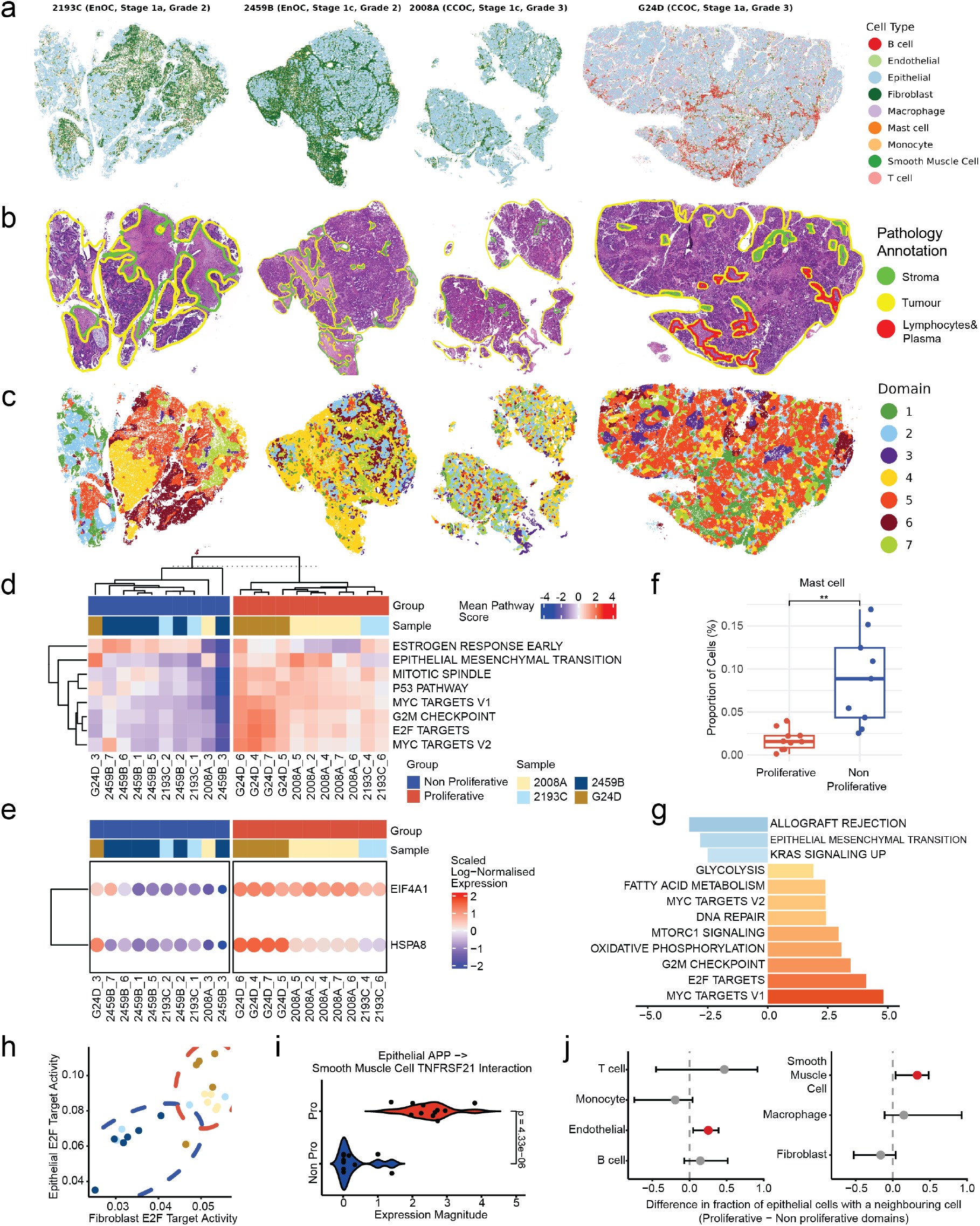
DOMINO resolves conserved proliferative programs across endometrioid and clear cell ovarian carcinomas. **a, b.** Spatially resolved maps from four archived tumours profiled with 10x Xenium, including two clear cell ovarian carcinomas (CCOC; 2008A, G24D) and two endometrioid ovarian carcinomas (EnOC; 2193C, 2459B). **c**,**d**. Increasing resolution to seven domains per sample revealed a conserved proliferation axis that stratified domains into proliferative and non-proliferative programs, supported by pathway activity clustering. **e**. Across all samples, tumour epithelial cells in proliferative domains showed significantly higher expression of FOXJ1, EIF4A1, and HSPA8 **f**. Relative abundance of mast cells within proliferative and non-proliferative domains across all tumours. Each point represents a single domain. **g**. Hallmark pathways significantly associated with fibroblasts residing within proliferative domains. The bar plot shows normalised enrichment scores (NES) from gene set enrichment analysis comparing fibroblasts from proliferative and non-proliferative domains. Positive NES values indicate enrichment in proliferative domains, whereas negative NES values indicate enrichment in non-proliferative domains. **h**. Correlation between E2F target activity in epithelial cells and fibroblasts across domains. Each point represents a single domain, and colours indicate individual tumours. Dashed ellipses denote proliferative and non-proliferative domains. **i**. Ligand–receptor analysis revealed significantly stronger epithelial–stromal communication in proliferative domains, highlighted by elevated APP–TNFRSF2 signalling between epithelial and smooth-muscle cells. **j**. Spatial proximity analysis between epithelial cells and neighbouring cell types from the same domain. The forest plot shows the difference in the fraction of epithelial cells with a neighbouring cell type within 100 μm in proliferative versus non-proliferative domains. Positive values indicate enrichment in proliferative domains. Error bars represent 95% confidence intervals and coloured points denote statistically significant differences after multiple testing correction.

To determine whether DOMINO domains captured distinct tumour states, we focused on tumour-enriched domains (>50% tumour cells). Hallmark pathway scoring analysis revealed the coexistence of proliferative and non-proliferative tumour states in three of the four tumours analysed (Fig. 3d). Sample 2459B lacked a proliferative state and was composed almost entirely of non-proliferative tumour cells (Fig. 3d). Compared to proliferative tumour cells, non-proliferative tumour cells showed reduced activity of proliferation-associated pathways, including MYC targets, together with increased estrogen signalling. Based on the dominant tumour states, tumour-enriched domains were subsequently classified as either proliferative or non-proliferative domains. Consistent across samples, *EIF4A1* and *HSPA8* are significantly upregulated in tumour cells from proliferative domains (Fig. 3e, Supplementary Fig. 25e, Supplementary Table 4).

To assess whether these findings generalised to independent datasets, we applied DOMINO to a recently published CCOC Xenium 5K spatial transcriptomic dataset generated by Le et al. ^32^. In contrast to our cohort, this tumour was less tumour-enriched, with tumour epithelial cells comprising only 33% of the 459,575 profiled cells, while fibroblasts accounted for 38% (Supplementary Figure 26a). Following segmentation into seven DOMINO domains, five domains (domains 2, 3, 5, 6, and 7) contained more than 15% tumour epithelial cells and were included in subsequent analyses (Supplementary Figure 26b). Hallmark pathway scoring revealed that tumour cells in domain 2 displayed markedly reduced activation of proliferation-associated pathways compared with tumour cells from the remaining tumour-containing domains (Supplementary Figure 26c). Consistent with our observations in the discovery cohort, \textit{EIF4A1} and \textit{HSPA8} were significantly upregulated in tumour cells from proliferative domains relative to tumour cells from the non-proliferative domain, supporting the existence of recurrent proliferative tumour states across independent CCOC spatial transcriptomic datasets (Supplementary Figure 26d). Comparable validation in an independent EnOC ST dataset was not possible because, to our knowledge, no publicly available EnOC ST datasets currently exist. Together, these results identify conserved epithelial programs associated with proliferative and non-proliferative tumour states, which were observed across multiple EAOC tumours and independently validated in an external CCOC spatial transcriptomic dataset.

Besides epithelial transcriptional differences revealed by DOMINO, the surrounding TME exhibited coordinated changes associated with tumour state. By comparing the cellular composition between proliferative and non-proliferative domains, we found that proliferative domains were characterised by reduced mast cell abundance across the cohort (Fig. **??**f). Owing to the presence of only a single non-proliferative domain in the independent CCOC dataset from Le et al. ^32^, formal statistical comparisons of cellular composition could not be performed. Nevertheless, the non-proliferative domain (domain 2) contained a higher proportion of mast cells than any of the tumour-enriched proliferative domains (Supplementary Figure 26f), consistent with the pattern observed in our discovery cohort.

Fibroblasts residing within proliferative domains displayed increased activity of proliferation-related pathways together with reduced allograft rejection, epithelial-mesenchymal transition and KRAS signalling (Fig. 3g). Notably, E2F target activity in fibroblasts strongly correlated with E2F target activity in neighbouring epithelial cells across domains, suggesting coordinated epithelial and stromal proliferation programs (Fig. 3h). Ligand receptor analysis further identified significantly increased signalling between epithelial *APP* and smooth muscle cell *TNFRSF21* in proliferative domains (Fig. 3i). Consistent with these signalling differences, epithelial cells within proliferative domains were significantly more likely to have neighbouring endothelial and smooth muscle cells within a 100 *μ*m radius around them than epithelial cells in non-proliferative domains (Fig. 3j), indicating preferential spatial association with vascular and stromal components. Collectively, these findings demonstrate that DOMINO domains capture coordinated epithelial and microenvironmental programs associated with tumour proliferation, encompassing altered cellular composition, stromal transcriptional states, intercellular signalling and spatial organisation.

### DOMINO identifies subtype-specific and tumour-excluded spatial domains shared across EAOCs

We next applied DOMINO to all four samples simultaneously, integrating data across the two Xenium slides to identify seven spatial domains shared across the cohort. The objective of this integrative analysis differs from the sample-specific analyses presented above. Whereas analysing each tumour independently enabled the identification of within-tumour spatial heterogeneity, the integrated analysis aimed to uncover spatial domains that are tumour-specific, subtype-specific, or shared across samples. DOMINO identified seven integrated spatial domains and revealed extensive transcriptional divergence between CCOC and EnOC (Fig. 4a, b, Supplementary Fi. 26a). This analysis demonstrates the scalability of DOMINO, which was able to integrate and analyse more than 641,000 cells across four tumours in a single framework. An alluvial plot linking tumour subtypes, integrated domains, and cell types demonstrated strong subtype-specific segregation (Fig. 4b). Domains 1 and 6 were composed almost exclusively of cells from the two EnOC samples, whereas domain 3 was predominantly derived from the two CCOC samples. Notably, this segregation was observed across both tumour and stromal populations, indicating that the two EAOC subtypes are characterised by distinct multicellular ecosystems rather than differences confined to the epithelial compartment. Comparison of tumour cells from the EnOC-associated domains (domains 1 and 6) and the CCOC-associated domain (domain 3) identified robust subtype-specific marker genes (Fig. 4c, Supplementary Table 5). These included significant upregulation of *CTNNB1*, a gene frequently mutated in EnOC ^33;34^, and elevated expression of *STAT3* in CCOC, consistent with activation of the *IL6* –*STAT3* signalling axis characteristic of this subtype ^35;36^. Together, these findings further support the presence of distinct epithelial programs underlying the two EAOC subtypes. In addition to the subtype-associated domains, DOMINO identified a distinct stromal domain (domain 4) composed almost exclusively of non-malignant cells and represented across all four tumours, irrespective of histological subtype(Fig. 4b,d). Spatially, domain 4 was largely tumour-excluded, with cells assigned to this domain located significantly further from tumour cells than cells in other domains (Fig. 4e, Supplementary Fig. 26a). Importantly, this tumour-excluded stromal state was only revealed through the integration of spatial information and transcriptional profiles, as these cells would otherwise appear as a stromal population shared between CCOC and EnOC, with no indication that their shared transcriptional profile reflects a common tumour-excluded spatial niche. The emergence of a dedicated stromal domain suggests that stromal cells residing in tumour-excluded regions adopt transcriptional states that are distinct from stromal cells co-localising with tumour cells. Consistent with this observation, several stromal cell populations separated according to their domain of residence, forming distinct clusters corresponding to CCOC-associated domains (domain 3), EnOC-associated domains (domains 1 and 6), and the tumour-excluded stromal domain (domain 4) (Fig. 4f, Supplementary Fig. 26b). This pattern was particularly evident in fibroblasts, which displayed clear transcriptional divergence across all three spatial contexts, suggesting that fibroblast state is shaped by both tumour subtype and spatial proximity to tumour cells (Fig. 4f). Additionally, fibroblasts in tumour-excluded regions exhibited reduced activity of proliferation-related pathways and increased inflammatory signalling (Fig. 4g). Together, these results reveal that EAOCs are organised into distinct multicellular ecosystems. They comprise subtype-associated tumour and stromal states as well as tumour-excluded stromal domains, highlighting the ability of DOMINO to uncover spatially defined cellular programs from complex ST data of ovarian cancer.

**Figure 4:**
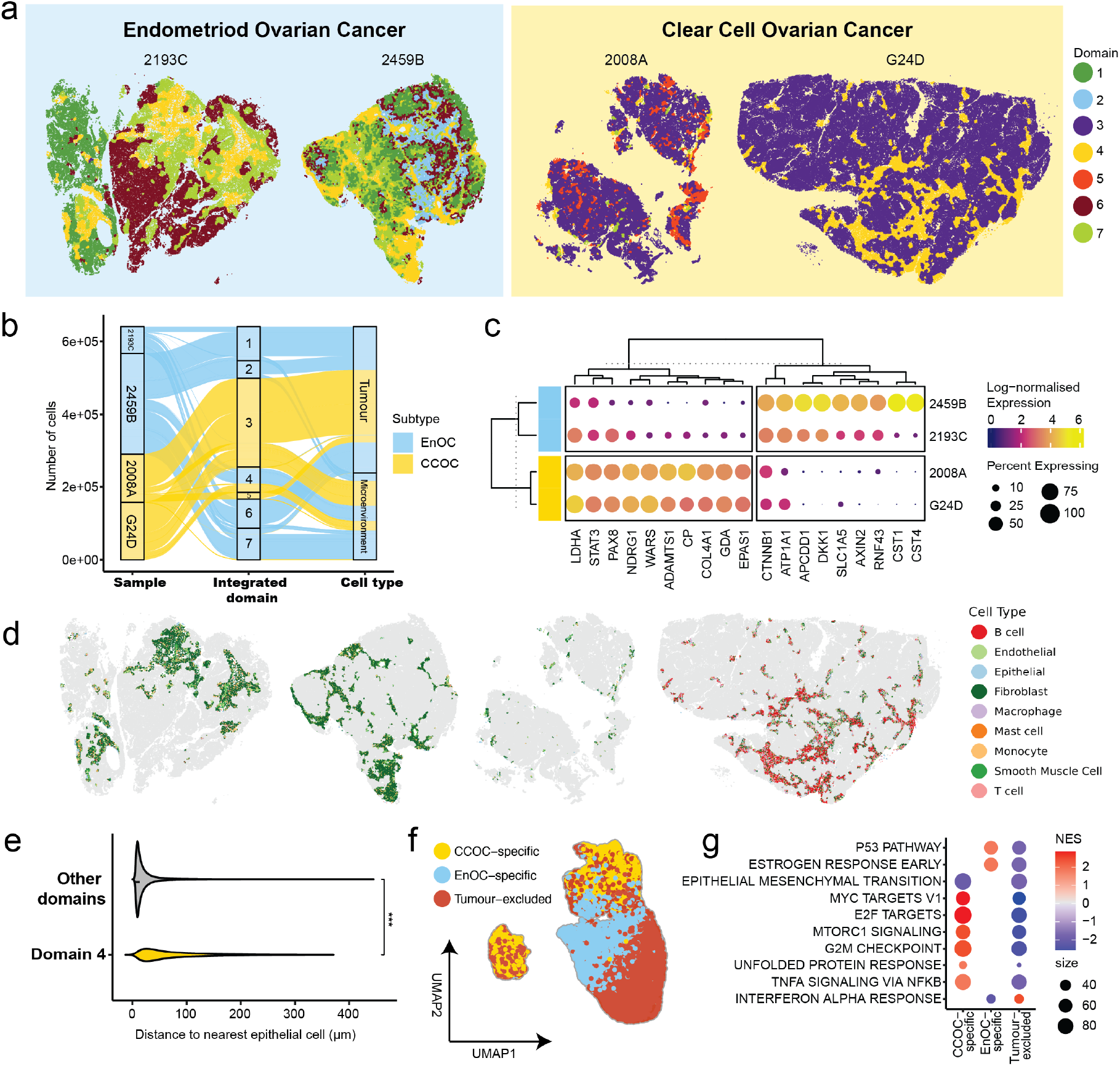
Integrative analysis via DOMINO reveals distinct spatial domains associated with EAOC subtype identity. **a.** Spatial distribution of the seven integrated DOMINO domains identified across all four tumours. Cells are coloured by integrated domain assignment. **b**. Alluvial plot linking tumour subtype, integrated domain, and cell type, demonstrating subtype-specific segregation of both malignant and stromal populations. **c**. Dot plot showing marker genes distinguishing tumour cells from EnOC-associated domains (domains 1 and 6) and the CCOC-associated domain (domain 3). Markers were required to be expressed in at least 70% of epithelial cells and exhibit | log_2_ FC| > 2. **d**. Spatial localisation of cells assigned to the tumour-excluded stromal domain (domain 4), coloured by cell type. **e**. Distance from cells in domain 4 and cells from all other domains to the nearest epithelial cell, demonstrating the tumour-excluded nature of domain 4. **f**. UMAP embedding of fibroblasts coloured according to their domain of residence, showing segregation of fibroblasts from CCOC-associated, EnOC-associated, and tumour-excluded domains. **g**. Hallmark pathways were significantly enriched in fibroblasts from CCOC-associated, EnOC-associated, and tumour-excluded domains. Tumour-excluded fibroblasts display reduced proliferation-associated pathways and increased inflammatory signalling relative to fibroblasts residing within tumour-associated domains.

## Discussion

Reconstructing domains from spatial transcriptomic (ST) maps allows to understand how cells organise, relate to each other and function in tissues. The spatial arrangement of cells underpins processes such as cell signalling, cell differentiation, and disease progression, with distinct tissue domains reflecting microenvironment and functional compartments. Here, we developed DOMINO, a diffusion-optimised graph contrastive learning framework designed for robust and scalable spatial domain identification. Through comprehensive benchmarking across diverse ST datasets and it application to a discovery cohort of ovarian cancer, DOMINO consistently scale up, allowing to process all dataset and outperformed existing methods in recovering biologically meaningful domains with better-defined boundaries, demonstrating both superior accuracy and scalability. These results should be interpreted in the context of an inherent limitation of spatial domain benchmarking: reference annotations are often the best available approximation of tissue organisation rather than absolute ground truth. The annotations used across benchmark datasets were derived from different sources, including pathological review, anatomical labels, or annotations provided by the original studies, and therefore vary in their biological granularity. For this reason, we evaluated DOMINO using multiple complementary quantitative metrics and qualitative spatial assessments. We believe this provides a more balanced assessment of domain concordance, while acknowledging that perfect agreement with reference labels is not expected across all datasets.

DOMINO builds upon the success of the graph-based contrastive learning model architecture ^37^. Graph-based neural networks naturally fit ST data and have been used to develop methods, such as GraphST ^17^ and Stagate ^16^. Moreover, several recent spatial-omics methods have incorporated diffusion-related ideas, the mechanism and objective of diffusion in DOMINO are fundamentally different. DiffusionST^38^ uses diffusion in the sense of deep generative diffusion modelling, which is primarily designed for representation enhancement, data generation or completion. This differs from the graph diffusion convolution used in DOMINO, which is a graph-structured signal propagation operator. DOMINO’s approach is also distinct from SDUCL, where signal diffusion is used to define local cellular microenvironments and the global representation is obtained by averaging node embeddings within those microenvironments ^39^. DOMINO instead uses the diffusion graph itself as an augmented structural view in a multi-view graph contrastive learning framework. By jointly contrasting the original graph, diffusion-enhanced graph and shuffled graph views, DOMINO derives representations that integrate local spatial-expression similarity, higher-order tissue topology and global graph-level structure. Thus, DOMINO is not a simple combination of existing diffusion and contrastive learning components but introduces a distinct diffusion-based graph augmentation strategy tailored to spatial domain identification. This design enables more coherent and stable domain recovery across heterogeneous tissue architectures. From an implementation perspective, DOMINO’s scalability arises from this graph-based design together with practical memory optimisation, rather than from keeping all data structures sparse throughout the method. In the current implementation, the neighbourhood and diffusion graphs are stored and processed sparsely, whereas the expression matrix is represented densely during model training. Scalability is therefore supported by sparse graph operations and chunked reconstruction loss computation.

While expert histopathological annotation remains foundational to cancer tissue interpretation, its scope can be challenged or augmented by the increasing resolution and molecular depth of modern ST datasets. Expert labelling defines regions at the morphological level, whereas ST captures molecular heterogeneity at near single-cell resolution, details that can reveal gradients and transitions not readily visible under the microscope. As data size and resolution continue to grow, manual annotation becomes increasingly time-consuming and difficult to scale. In this setting, DOMINO complements pathologist-defined regions by providing an unsupervised data-driven framework that infers molecular-coherent domains directly from ST data. Importantly, its tunable resolution allows users to explore tissue architecture at multiple scales: from broad domains that align closely with histopathological annotations to high-resolution domains that capture finer molecular granularity within morphologically similar regions.

Endometriosis-associated ovarian cancers (EAOCs), comprising endometrioid (EnOC) and clear cell ovarian carcinomas (CCOC), are clinically and molecularly distinct from high-grade serous ovarian cancer (HGSOC). Although often diagnosed in the early stages, they respond poorly to platinum-based therapy and have limited treatment options once advanced ^28^, highlighting the need for deeper molecular characterisation and new therapeutic avenues. Despite their clinical relevance, EAOCs the interactions between their epithelial compartments and the surrounding microenvironment are poorly defined. In this study, we present the first high-resolution, high-plex spatial transcriptomic comparison of the two major EAOC subtypes, EnOC and CCOC, providing a unified view of their epithelial states, stromal organisation, and multicellular ecosystems. By applying DOMINO to ST data from EAOCs, we were able to address these gaps and resolve tumour architectures that cannot be captured by histology, bulk profiling, or conventional single-cell analyses.

A major finding of this study is that EAOCs are organised into spatially distinct multicellular ecosystems (*i*.*e*., domains) that extend beyond the epithelial compartment. While previous studies have established substantial molecular and clinical differences between EnOC and CCOC ^23;26^, our integrated analysis demonstrates that these differences are also reflected in their spatial organisation. EnOC- and CCOC-associated domains contained distinct epithelial programs together with transcriptionally divergent fibroblast populations, indicating that the biological differences between these subtypes are not restricted to tumour cells but encompass the broader tissue microenvironment. These findings suggest that the divergent clinical behaviour of EnOC and CCOC may arise, at least in part, from subtype-specific tumour-microenvironment interactions.

Beyond subtype-associated domains, we identified a tumour-excluded stromal niche that was conserved across tumours. Cells assigned to this domain were spatially separated from tumour epithelium and exhibited transcriptional programs distinct from stromal cells co-localising with tumour cells. Fibroblasts within this niche displayed reduced proliferative activity and increased inflammatory signalling relative to fibroblasts residing within tumour-associated domains, indicating that spatial proximity to tumour cells is a major determinant of stromal cell state. This observation is consistent with accumulating evidence that fibroblast identity is highly plastic and can be shaped by local tumour-derived signals, giving rise to diverse cancer-associated fibroblast states within the tumour microenvironment ^40;41^. Importantly, this stromal domain emerged only when spatial information was incorporated into the analysis. Without spatial context, these cells would likely have been grouped with transcriptionally similar stromal populations despite occupying a distinct anatomical niche. Together, these findings highlight an important advantage of spatially informed domain detection and suggest that tumour-associated stromal states may arise through local reprogramming of resident stromal cells by neighbouring tumour cells.

In addition to subtype-specific and tumour-excluded ecosystems, we identified conserved proliferative and non-proliferative tumour states across multiple tumours. Proliferative domains were characterised by increased expression of *EIF4A1*, and *HSPA8*, together with enrichment of cell cycle-associated pathways, whereas non-proliferative domains displayed increased estrogen signalling. This is consistent with prior ovarian cancer studies implicating *EIF4A1* in translational control, including FXR1-mediated recruitment of the eIF4F complex to promote cMYC translation, and identifying *HSPA8* as a regulator of CLPP stability, mitophagy and cisplatin resistance in ovarian cancer cells ^42;43^. The recurrence of these states across multiple EAOC tumours, together with their independent validation in an external CCOC spatial transcriptomic dataset, suggests that they represent conserved epithelial programs rather than patient-specific features. Notably, these epithelial states were accompanied by coordinated changes in the surrounding microenvironment, including reduced mast cell abundance in proliferative domains, distinct fibroblast transcriptional states, altered vascular association, and domain-specific intercellular signalling. Although the role of mast cells in ovarian cancer remains incompletely understood, several studies have reported tumour-suppressive functions ^44^ and favourable clinical associations for mast cell infiltration ^45^, raising the possibility that mast cell depletion may contribute to the establishment of highly proliferative tumour microenvironments. Together, these observations support a model in which tumour proliferation is not solely a cell-intrinsic property but instead emerges within a broader spatially organised ecosystem involving epithelial, stromal, immune, and vascular components.

An important future direction will be to determine whether the spatial domains identified here are associated with clinical outcomes. The recurrent subtype-specific, proliferative, and tumour-excluded domains observed across tumours suggest that spatial architecture may provide prognostic information beyond conventional histopathological classification. However, such analyses remain challenging because both EnOC and CCOC are relatively uncommon ovarian cancer subtypes and are consequently underrepresented in existing spatial transcriptomic cohorts. Consistent with this limitation, we explored whether expression of *EIF4A1* and *HSPA8*, markers associated with proliferative domains, was associated with overall survival in publicly available ovarian cancer datasets. Owing to the limited representation of clear cell and endometrioid tumours and the small number of survival events, no significant associations were identified (Supplementary Fig. 26g, h). As larger datasets with linked clinical follow-up become available, it will be possible to evaluate whether specific spatial domains, multicellular ecosystems, or patterns of domain organisation are associated with recurrence, treatment response, or patient survival.

## Methods

### DOMINO

#### Data preprocessing within DOMINO

During the data preprocessing stage, the top 2000 most highly variable genes (HVGs) are selected by default, or the full gene panel is used if fewer than 2,000 genes are available. This step aims to reduce data dimensionality, minimise noise interference, and highlight genes exhibiting significant biological differences, thereby enhancing the accuracy and interpretability of subsequent analyses. Subsequently, normalisation is performed, followed by logarithmic transformation using the SCANPY package to make the data distribution closer to a normal distribution and stabilise the variance ^46^. Finally, feature scaling is performed to scale the expression values of each gene to a unit variance, further enhancing model training stability.

#### Graph construction

Gene expression and cell coordinates from spatial transcriptomics data were used to construct an undirected neighbourhood graph *G* = (*V, E*). In the graph *G*, **V** represents the set of cells/spots while **E** represents the set of edges between them. Gene expression information is used as node feature **F** while the spatial position of cells is used as node feature. BallTree is used to find the nearest neighbours, obtain the indices of each spot and its nearest neighbours, and construct the original adjacency matrix ^47^. The adjacency matrix is defined as **A** ∈ ℝ^*N* ×*N*^ with *N* denoting the number of cells/spots. The original unidirectional adjacency matrix is used as a mask for subsequent pooling operations, and a bidirectional symmetric adjacency matrix is generated for neighbourhood aggregation in graph neural networks (GNNs).

#### Graph diffusion enhancement

To generate the diffusion graph, we use graph diffusion convolution (GDC) to process the original graph. The adjacency matrix is transformed into a diffusion matrix, thereby achieving graph augmentation for the original graph. This process efficiently approximates the Personalised PageRank (PPR) matrix via the Arnoldi iteration method.

The formula for the overall diffusion process is as follows:

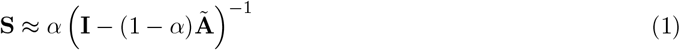

where **I** is the unit matrix, **Ã** is the symmetric normalised adjacency matrix, and *α* is the restart probability parameter to maintain the balance between local and global aspects. **S** is the approximate PPR diffusion matrix, which includes the PPR similarity between all pairs of nodes; that is, *S*_*i,j*_ represents the probability of a random walk starting from node *i* and reaching node *j*.

The specific steps for graph diffusion enhancement are as follows:

1. Add self-loops to the original adjacency matrix:

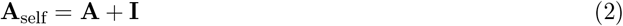

where **A**_self_ represents the adjacency matrix after adding self-loops. This step ensures that each node is connected to itself.
2. Perform symmetric normalisation on **A**_self_ to obtain the symmetric normalised adjacency matrix **Ã** :

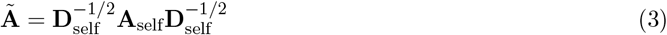

where **D**_self_ is the degree matrix after adding self-loops, defined as *D*_self,*i,i*_ =∑_*j*_*A*_self,*i,j*_.

During the diffusion computation stage, the Arnoldi iterative method (which accelerates convergence) is used for iterative initialisation, defined as:

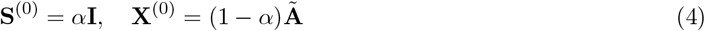

Subsequently, continuous iteration, update, and convergence checks are performed until the diffusion matrix is obtained. The iterative update steps are:

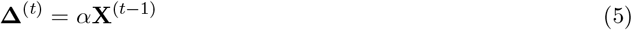

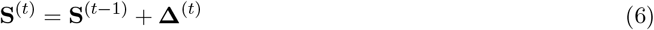

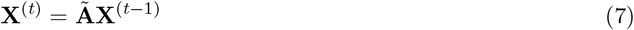

where **S**^(*t*)^ is the diffusion matrix after the *t*-th iteration, **X**^(*t*)^ is the intermediate matrix of the *t*-th iteration, and **Δ**^(*t*)^ is the increment matrix of the *t*-th iteration.

The convergence check condition is defined as:

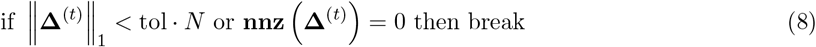

where tol is the convergence tolerance threshold, nnz(·) is the function for counting the number of non-zero elements in a matrix, and ∥ · ∥_1_ denotes the matrix 1-norm (sum of absolute values of all elements).

After obtaining the diffusion matrix, post-processing (top-*k* sparsification and row normalisation) is performed:

1. Top-*k* sparsification: Retain the *k* edges with the highest weights for each node *i*:

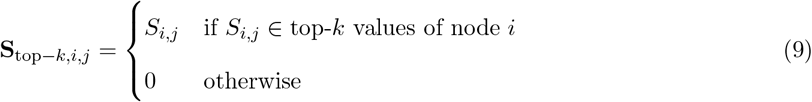

where *k* is the maximum number of edges per node, and **S**_top−*k*_ is the diffusion matrix after sparsification.
2. Row normalisation:

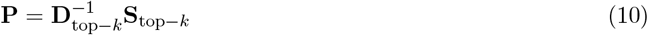

where **D**_top−*k*_ is the degree matrix of **S**_top−*k*_ (i.e., *D*_top−*k,i,i*_ = ∑_*j*_ *S*_top−*k,i,j*_), and **P** is the final diffusion matrix.

#### Multi-view graph contrastive learning

To enable the model to capture both local and global (diffusion) graphs, a self-supervised graph contrastive learning framework is constructed for model training. Specifically, graph contrastive learning is used as the basic graph encoder; the diffusion graph and the original graph are fed into two separate GCNs, followed by a shared projection head (a multi-layer perceptron (MLP) with two hidden layers and a Parametric Rectified Linear Unit (PReLU) non-linearity) to learn node representations for the two views. Subsequently, representations from the GCN outputs are fed into a pooling layer and the shared MLP to learn graph representations for the two views. Finally, a discriminator contrasts node representations from one view with graph representations from the other view (and vice versa) and scores their agreement. For constructing the graph contrastive loss function:

1. Define a binary label matrix **Y** ∈ ∑^*N* ×1^, where *Y*_*i*,1_ = 1 (positive samples) and *Y*_*i*,1_ = 0 (negative samples).
2. The graph contrastive loss has two components:

Component 1: The node representation **Z** (from GCN encoding of the original graph) and the graph representation **S** (from GCN encoding of the diffusion graph) form positive pairs; the shuffled node representation **Z**^′^ (from shuffled original graph features) and **S** form negative pairs. The loss is:

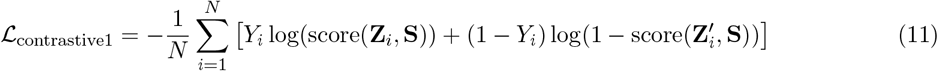

where score(**z, s**) = *σ*(**z**^*T*^ **s**), and *σ*(·) is the sigmoid function (discriminator score).

Component 2: The node representation **Z**_*d*_ (from GCN encoding of the diffusion graph) and the graph representation **S**_*o*_ (from GCN encoding of the original graph) form positive pairs; the shuffled node representation **Z**^′^ and **S**_*o*_ form negative pairs. The loss is:

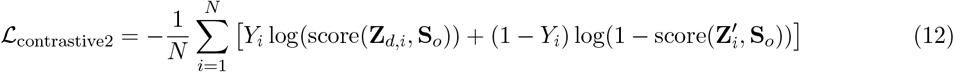

where score(**z, s**_*o*_) = *σ*(**z**^*T*^ **s**_*o*_).

#### Model training and loss function

The total loss consists of two parts: graph contrastive loss and feature reconstruction loss.

1. Feature reconstruction loss: Calculated by the mean square error (MSE) between the original node features **F** and the reconstructed features 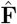 (output by the decoder from GCN-encoded node representations):

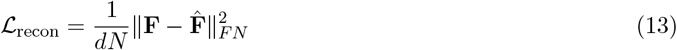

where *d* is the dimension of input node features, and ∥ · ∥_*F*_ is the Frobenius norm (square root of the sum of squares of all elements).
2. Total loss:

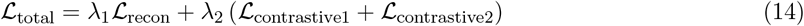

where *λ*_1_ and *λ*_2_ are weight hyperparameters for balancing the reconstruction task and graph contrastive learning.

#### Sparse representation and scalability

To avoid storing neighbourhood graphs and diffusion matrices as dense *N* × *N* arrays to improve scalability. We reformulated these structures as sparse coordinate tensors, retaining only non-zero entries and adding self-loops as a sparse diagonal. All downstream operations were engineered to rely on sparse matrix-vector multiplication rather than dense broadcasting, thereby reducing the memory complexity of DOMINO from *O*(*N* ^2^) to approximately *O*(*kN* ), where *k* denotes the average number of neighbours per cell.

This sparse reformulation substantially improved scalability in practice. On an A800 GPU (80 GB of device memory) with 100GB of RAM, DOMINO successfully processed datasets containing up to 600,000 cells with 2000 highly variable genes using default hyperparameters. This scalability is particularly important as advances in spatial transcriptomics technologies continue to increase the number of captured cells, positioning DOMINO as a practical and robust choice for large-scale spatial analyses.

#### Spatial domain identification

After model training, the reconstructed feature matrix 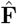 is input into a clustering algorithm (mclust by default, Leiden and Louvain are optional) ^48–50^ to assign spatial domain labels to each spot, completing spatial domain identification.

To further improve clustering accuracy, we also include a cluster refinement step by including a post-processing optimiser. The optimiser examines the cluster composition in the local neighbourhood of each spot and refines initial clustering labels via an ensemble system to produce spatially continuous and consistent results. For large-scale data, a block-based strategy is used to compute the spatial distance matrix, reducing memory consumption.

#### Multi-sample domain identification

For multi-sample spatial domain identification, samples were analysed jointly after harmonising their feature and coordinate spaces. Each sample was first restricted to the set of genes shared across all samples, ensuring that the concatenated object was represented by a common expression feature space. Spatial coordinates were then transformed sample-wise by shifting each sample to a common origin and translating samples along the x-axis with a fixed inter-sample gap. This preserved within-sample spatial organisation while preventing artificial nearest-neighbour connections between independent samples during graph construction.

The transformed samples were concatenated into a single object. DOMINO was trained directly on this joint object to obtain a shared embedding across samples. In this study, to reduce sample-specific effects, Harmony correction ^31^ was applied to the DOMINO embedding using the sample label as the batch covariate. The corrected embedding was then used for downstream spatial domain identification with mclust, followed by local label refinement as described above. This strategy enabled domains to be identified jointly across samples while preserving sample-specific spatial topology.

### Benchmarking analysis

#### Domain identification benchmarking

To evaluate the performance between predicted spatial domains of each method against ground truth annotations, we used multiple metrics including Adjusted Rand Index (ARI), Normalised Mutual Information (NMI), Adjusted Mutual Information (AMI), Normalised Variation of Information (NVI), Fowlkes–Mallows Index (FMI) and Purity. ARI, AMI, NMI and NVI were computed with the `aricode` package ^51^, while FMI was obtained either from `dendextend`^52^.

Purity was computed directly using the following definition. For a set of reference labels 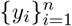 and predicted labels 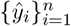, let *C* denote the set of predicted clusters and *T* the set of true classes. Construct the contingency table *N*_*t,c*_ counting points of true class *t* ∈ *T* assigned to cluster *c* ∈ *C*. The clustering purity is

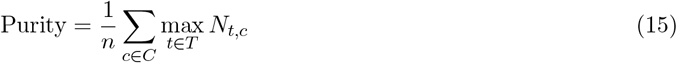

that is, the fraction of observations that belong to the majority true class within their predicted cluster.

To summarise performance across heterogeneous metrics, we first scaled ARI to [0, 1] by ARI_01_ = (ARI + 1)*/*2, inverted NVI to NVI_*i*_ = 1 − NVI (so that larger values always indicate better performance), and applied *z*-score normalisation per metric across methods. The mean *z*-score defined an overall score for each method.

#### Domain boundary detection benchmarking

To complement clustering-based benchmarking, we designed two graph-based metrics to quantify the performance of identifying spatial domain boundaries: *Boundary Purity* (BP) and *Boundary Consistency* (BC). Both metrics operate on a *k*-nearest-neighbour (kNN) graph constructed from the spatial positions of spots or cells, where vertices represent observations and edges connect each vertex to its *k* nearest neighbours in Euclidean space. Let *G* = (*V, E*) denote this graph, with |*V* | = *n* vertices and labels 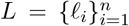 corresponding to predicted domains.

Boundary Purity measures the degree of label mixing at domain boundaries. For each vertex *i* ∈ *V*, we define its one-hop neighbourhood *N* (*i*) as the set of adjacent vertices in *G*. The local purity is

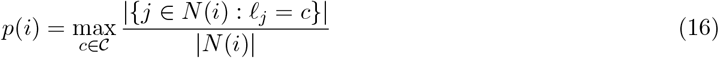

where *C* is the set of unique domain labels. Vertices located on a boundary are identified as

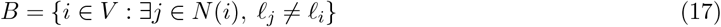

The Boundary Purity for the whole dataset is the mean purity over all boundary vertices:

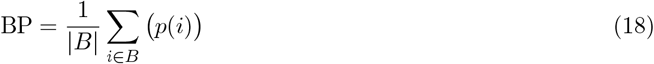

High BP indicates that cells along domain interfaces remain locally homogeneous, reflecting crisp borders, whereas low BP reflects label mixing and diffuse boundaries.

Boundary Consistency evaluates the stability of regions flanking a boundary. Let *E*_*b*_ = {(*u, v*) ∈ *E* : *ℓ*_*u*_≠ *ℓ*_*v*_} denote the set of boundary edges. For each endpoint *u* of a boundary edge (*u, v*), we consider its *h*-hop neighbourhood excluding *v*,

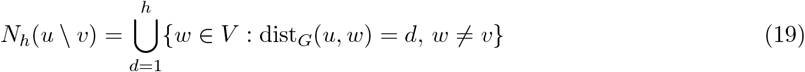

and compute its purity *p*_*h*_(*u* \ *v*) as previous.

Each boundary edge receives a score equal to the average purity on both sides:

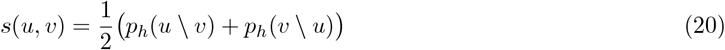

and the Boundary Consistency is the mean over all boundary edges,

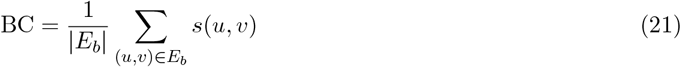

High BC indicates that, once across a boundary, labels quickly converge to a single domain, capturing the stability and clarity of predicted boundaries.

We implemented these two metrics using the RANN package ^53^ for efficient kNN search and `igraph`^54^ for graph operations. Unless otherwise specified, we used *k* = 6 to approximate hexagonal neighbourhoods typical of spot-based assays and set *h* = 2 when calculating BC.

For visualisation and comparison across datasets, BP and BC were normalised relative to the corresponding ground-truth annotation within each dataset. This dataset-wise normalisation accounts for differences in intrinsic boundary complexity across tissues and platforms. For both metrics, values closer to 1 indicate boundary properties more similar to the annotated ground truth, whereas values deviating from 1 indicate boundaries that are either less locally coherent or overly simplified relative to the reference annotation.

#### Spatial data simulation for scalability benchmark

To assess scalability, we generated *in silico* spatial transcriptomics datasets of increasing sizes using a simple simulation pipeline. Gene expression counts were simulated as sparse matrices, where each cell contained a fixed fraction (0.05 by default) of non-zero genes with values drawn from a negative binomial distribution to mimic zero-inflated overdispersed scRNA-seq–like counts. Spatial coordinates were independently sampled from a uniform two-dimensional grid to create a neutral spatial layout without biological structure. The simulated counts and coordinates were assembled into AnnData objects and saved in compressed HDF5 format, enabling efficient creation of large-scale datasets for benchmarking runtime and memory performance.

### Seeding sensitivity analysis

#### Boundary and region sensitivity analysis

Boundary and region sensitivity were evaluated on a spatial k-nearest-neighbour graph constructed from the spot or cell coordinates of each dataset using *k* = 6. For each labelling, boundary nodes were defined by local label heterogeneity: a node was considered a boundary node if its neighbours contained at least two distinct labels and the second most abundant label represented at least 20% of the neighbourhood. Boundaries were identified separately for the ground-truth annotations and for each method. Boundary precision was defined as the fraction of predicted boundary nodes located within one graph hop of a ground-truth boundary node, and boundary recall as the fraction of ground-truth boundary nodes located within one graph hop of a predicted boundary node. Boundary F1 was calculated as the harmonic mean of boundary precision and recall.

To assess recovery of fine-scale annotated structures, each ground-truth label was decomposed into spatially connected components on the same graph, with each component treated as an individual annotated region. Regions were ranked by size within each dataset and assigned to small, medium or large bins using tertiles. For each annotated region, the predicted domain with the largest overlap was selected as its best match. Region recovery was quantified using best-match recall, defined as the overlap divided by the size of the annotated region, and best-match intersection-over-union (IoU), defined as the overlap divided by the union of the annotated region and the matched predicted domain. Recall, therefore, measures how completely an annotated region is recovered, whereas IoU additionally penalises over-merged predictions that absorb neighbouring regions.

### Ovarian tumour Xenium data

#### Sample collection and consent

The ovarian tumour samples used in this study were collected under approvals from the Central Adelaide Local Health Network Human Research Ethics Committee (HREC/18/CALHN/811, HREC/14/RAH/13, #19751) during 2016 to 2025. Formalin-fixed paraffin-embedded (FFPE), treatment-naive primary ovarian tumours from 4 patients diagnosed with stage I (grade 2/3) (Supplementary Table 3) were included in this study. To ensure the RNA quality of candidate FFPE blocks, 15 µm scrolls were sectioned from each block to measure DV200. Only blocks with DV200 ≥30% were selected for Xenium profiling.

#### Xenium spatial transcriptomic profiling

FFPE tissue sections of four archived tumours were prepared at the University of Adelaide Histology Service. For each case, a 5 µm section was cut and mounted, with two tumour sections placed on each of two Xenium slides (two sections per slide) (10x Genomics, PN-3000941). Slides underwent deparaffinization and decrosslinking according to the Xenium In Situ for FFPE Deparaffinization and Decrosslinking protocol (CG000580 Rev E, 10x Genomics). In situ gene expression was then profiled on the 10x Genomics Xenium Analyser using the Human Gene Expression 5K Prime probe set (10x Genomics, PN-1000724), following the Xenium Prime In Situ Gene Expression with Optional Cell Segmentation User Guide (CG000760 Rev C, 10x Genomics) and Xenium Analyser User Guide (CG000584 Rev F, 10x Genomics) without deviations. Xenium runs, and primary data generation was performed by the South Australian Genomics Centre (SAGC) using 10X’s default settings (Xenium instrument software version 3.4.1.0). Cell segmentation and probe ligation counting were carried out with the Xenium onboard analysis pipeline (version 3.3.0.1), and the outputs were exported from Xenium Analyser for downstream analyses.

#### Histopathological assessment

Haematoxylin and eosin (H&E) staining was performed on 5 µm sections consecutive to those profiled by Xenium. H&E staining was performed using the Dako Coverstainer automated stainer (Agilent Technologies) following the manufacturer’s protocol. Slides were then imaged using the Zeiss NanoZoomer 2.0-HT. A specialist gynecological pathologist reviewed the H&E slides using *Aperio ImageScope* to confirm tumour histological subtypes and delineate tumour and non-tumour regions.

#### Per-sample domain analysis

The per-cell feature count matrix, spatial coordinate file, and per-transcript table were imported into R as a `SpatialExperiment`^55^ object. Low-quality cells were removed on a per-sample basis if either the library size or the number of uniquely detected features was below (median 3×MAD) across all cells, using `scuttle`^56^. The remaining cells from all four tumours were then merged. Dimension reduction and clustering were performed using `Seurat`^57^, based on the top 2000 HVGs, the top 30 principal components, and the Leiden algorithm ^58^ (resolution 0.8).

Cell type labels were assigned using `SingleR`^59^ against three independent references: the Human Primary Cell Atlas (HPCA) ^60^, an integrated ovarian tumour scRNA-seq reference from Guimarães *et al*. ^61^, and scRNA-seq data from four ovarian tumours (three HGSOC and one endometrioid) from Regner *et al*. ^62^. For HPCA, only cell types expected in ovarian tumours were retained. Predictions across the three references were compared (Supplementary Fig. 27), and canonical marker expression was inspected to derive a consensus annotation.

DOMINO was applied to each tumour separately using the top 2000 HVGs identified from the merged dataset. Cells were scored against Hallmark gene sets ^63^ (retrieved via `msigdbr`^64^) using `UCell`^65^. Marker genes distinguishing proliferative and non-proliferative tumour states were detected using `FindMarkers` in each sample (min.pct = 0.8). Differential expression results in each tumour are available as Supplementary Table 4. Expression of genes significantly up-regulated (p.adj ¡ 0.05, log fold change > 0) across all samples was visualised using `ComplexHeatmap` ^66^. Differences in proportions of cell type between domains were tested using `propeller`^67^. Differentially expressed genes between fibroblasts residing in proliferative and non-proliferative domains were identified using `FindMarkers`, and significantly enriched gene sets were determined using `fgsea`^68^. Ligand–receptor inference was performed using Connectome ^69^ via the LIANA+ ^70^ wrapper within each domain in each tumour. Differences in ligand–receptor interaction activities between proliferative and non-proliferative domains were assessed using the Wilcoxon rank-sum test. To quantify local cellular neighbourhoods, the nearest neighbouring cell of each non-epithelial cell type was identified for every epithelial cell within the same sample and spatial domain using Euclidean distances calculated from spatial coordinates. A neighbouring cell was defined as being located within 100 µm of an epithelial cell. For each cell type and domain, the fraction of epithelial cells with at least one neighbouring cell within this radius was calculated. Differences between proliferative and non-proliferative domains were assessed using the Wilcoxon rank sum test, and effect sizes with 95% confidence intervals were estimated by bootstrap resampling (1,000 iterations) of the median difference in epithelial neighbour fractions between domain groups.

#### Cross-sample integrative domain analysis

Harmony was applied to DOMINO feature embeddings obtained following contrastive learning, and joint clustering was subsequently performed across all four EAOC samples to identify seven integrative spatial domains. Since each slide contained one clear cell and one endometrioid tumour, the slide was used as the batch correction variable to minimise technical variation while preserving biological differences between samples. Subtype-specific domains were defined based on the cellular composition of each domain. Marker genes distinguishing tumour epithelial cells from subtype-specific domains were identified using `FindMarkers`. Robust markers were defined as genes expressed in at least 70% of epithelial cells within one domain and exhibiting an absolute log_2_ fold change greater than 2 (| log_2_ FC| > 2). Differences in the distance of cells to the nearest epithelial cell between tumour-excluded and other domains were assessed using the Wilcoxon rank-sum test. Differential expression analysis between fibroblasts from different domain types was performed using `FindMarkers`, followed by gene set enrichment analysis using `fgsea`^68^.

## Data availability

Spatial transcriptomics data used in the benchmarking experiment of this study are all publicly available. The 10X Xenium datasets were retrieved from the 10X database at https://www.10xgenomics.com/datasets/ffpe-human-breast-with-custom-add-on-panel-1-standard (breast cancer) and https://www.10xgenomics.com/datasets/fresh-frozen-mouse-brain-replicates-1-standard (mouse brain). The CosMx non-small cell lung cancer (NSCLC) data were sourced from the official Nanostring website at https://brukerspatialbiology.com/products/cosmx-spatial-molecular-imager/ffpe-dataset/nsclc-ffpe-dataset/. The Stereoseq mouse brain and embryo data are downloaded from Chen *et al*. ^71^. The MERFISH mouse colon data were obtained from https://doi.org/10.5061/dryad.rjdfn2zh3^72^. The Visium DLPFC data is available from Maynard *et al*. ^73^. The Visium human breast cancer (https://www.10xgenomics.com/datasets/human-breast-cancer-block-a-section-1-1-standard-1-1-0) and mouse brain anterior data (https://www.10xgenomics.com/datasets/mouse-brain-serial-section-1-sagittal-anterior-1-standard-1-1-0) are from the 10X Visium demo database. The 10X Xenium ovarian cancer generated in this study can be accessed in Zenodo (doi: 10.5281/zenodo.17888636). Cell annotations that were used in this study were obtained from the original datasets where available; otherwise, pathological annotations from Bhuva *et al*.. ^74^ were used.

## Code availability

The source code of DOMINO is publicly available at the GitHub repository: https://github.com/ABILiLab/DOMINO. Analysis code used in this manuscript is available at the GitHub repository: https://github.com/ABILiLab/DOMINO_paper_code.

## Acknowledgments

This work was supported by the National Key Research and Development Program of China [2022YFF1000100]; the National Natural Science Foundation of China [62202388]; and the Qin Chuangyuan Innovation and Entrepreneurship Talent Project [QCYRCXM-2022–230]. F.L. was supported by the Australian National Health and Medical Research Council Investigator grants [2041439]. J.P. was supported by the Australian National Health and Medical Research Council Investigator grants [2035021]. The South Australian immunoGENomics Cancer Institute (SAiGENCI) received grant funding from the Australian Government. We thank Yvonne Ciuk from Histology Services at the Adelaide University for her assistance with Xenium slide preparation and Haematoxylin and eosin (H&E) staining.

## Contributions

F.L. and N.L. conceptualised and supervised the project with help from J.P.. P.J. designed the model with feedback from F.L. and N.L.. P.J. and N.L. developed the DOMINO software with feedback from F.L. and N.W.L.. P.J. and N.L. led the benchmarking analysis with feedback from J.P., N.W.L., and F.L.. X.G. and C.W. helped with the software development and benchmarking experiments. N.W.L., N.L., and P.J. wrote the manuscript with feedback from J.P. and F.L.. N.W.L. led the ovarian cancer data analysis with input from J.P. and N.L.. N.W.L., N.L., Z.R., P.J., X.G., M.M., and T.Z. contributed to figure design and generation. N.W.L., T.Z., and R.M. annotated and interpreted the ovarian cancer datasets. S.M. and E.W. helped with the 10X Xenium experiments and the H&E scan to generate the ovarian cancer spatial transcriptomic data. M.O., C.R., and N.A.L. performed tissue collection, archiving and selection of the ovarian cancer tissues used in this study.

### Corresponding author

Correspondence to Fuyi Li, Ning Liu, and Jose M Polo.

## Ethics declarations

All human tissue samples were obtained in accordance with institutional ethical guidelines and approved human research ethics protocols. Tumour samples were collected under approvals from the Central Adelaide Local Health Network Human Research Ethics Committee (HREC/18/CALHN/811, HREC/14/RAH/13, #19751). Informed consent was obtained from all participants where applicable, and all procedures were conducted in compliance with the National Statement on Ethical Conduct in Human Research (NHMRC, Australia).

## Additional information

## Supplementary information

