## supplementary information for "DOMINO: graph diffusion learning identifies spatial domain structures with enhanced accuracy and scalability"

### 1 Supplementary Figures

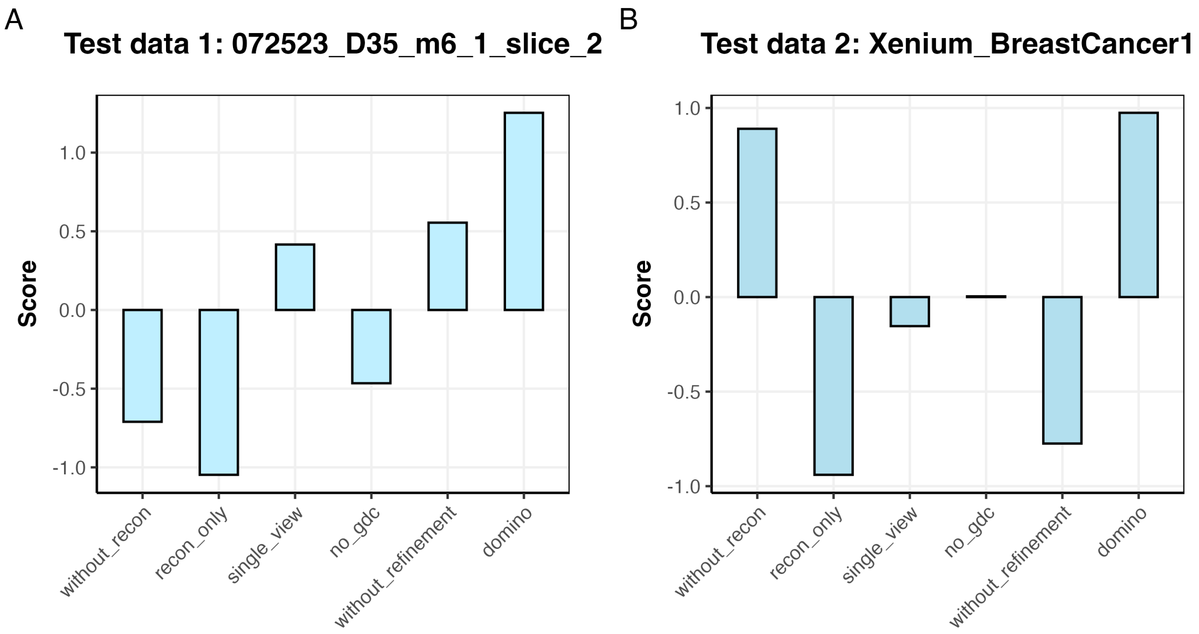


**Supplementary Figure 1.** Ablation analysis of DOMINO. To assess the contribution of each component of DOMINO, we compared the full model with five ablation settings. **Ablation study 1** used a reconstruction-only baseline comprising the original graph, a GCN encoder, and a decoder, trained solely with feature reconstruction loss and without graph contrastive learning. **Ablation study 2** evaluated a single-view graph contrastive model, in which node embeddings and pooled graph embeddings were contrasted within the same graph view, with negative samples generated from shuffled node features. **Ablation study 3** replaced the graph diffusion view with a feature-masking augmented view, in which a subset of node features was randomly set to zero, to test whether the graph diffusion view provides advantages over a generic augmentation strategy. **Ablation study 4** removed the feature reconstruction loss and retained only the two graph contrastive losses during training. **Ablation study 5** removed the post-clustering refinement step from the full DOMINO pipeline to evaluate whether performance and boundary quality depend primarily on representation learning rather than downstream label refinement. The analysis was performed on two downsampled MERFISH slices, *072523_D35_m6_1_slice_2* and *072523_D35_m11_1_slice_2*. Evaluation metrics were calculated as described in the **Methods** section.

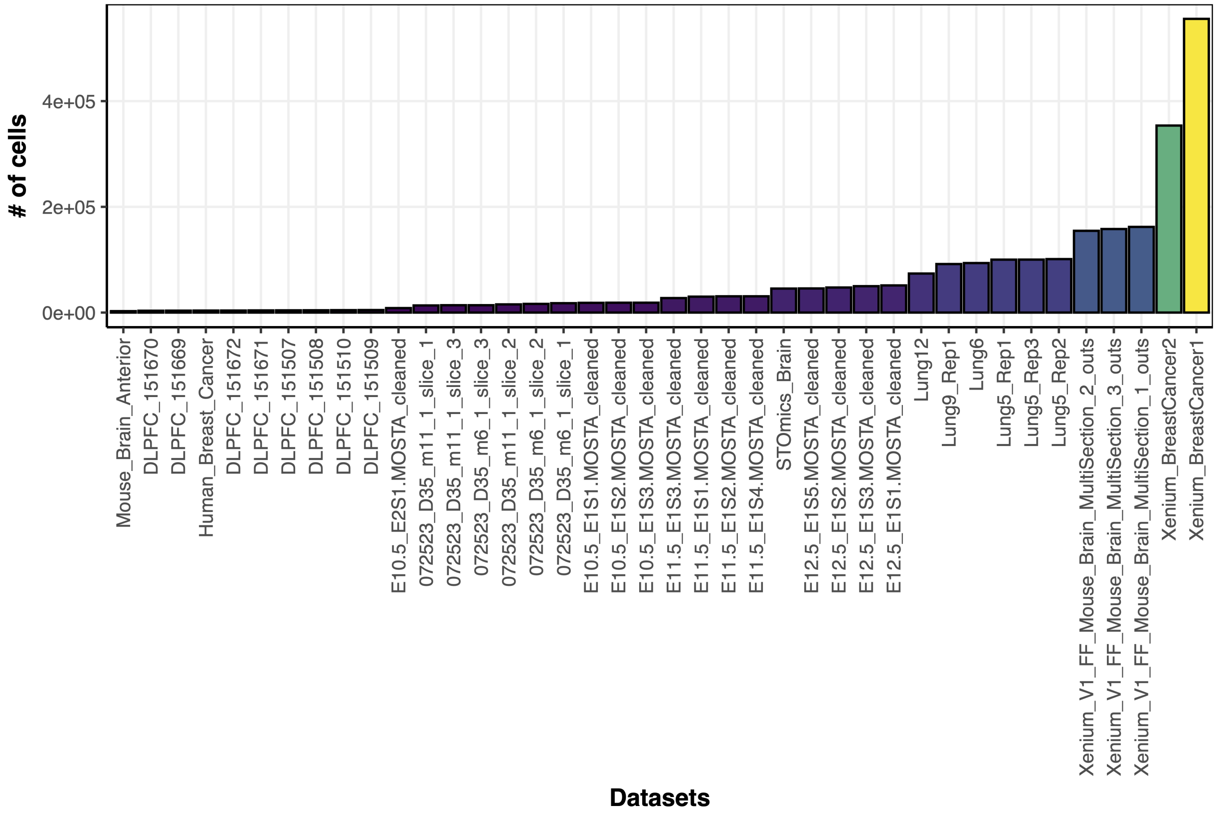


**Supplementary Figure 2.** Cell counts of spatial transcriptomics datasets that are used in benchmarking analysis.

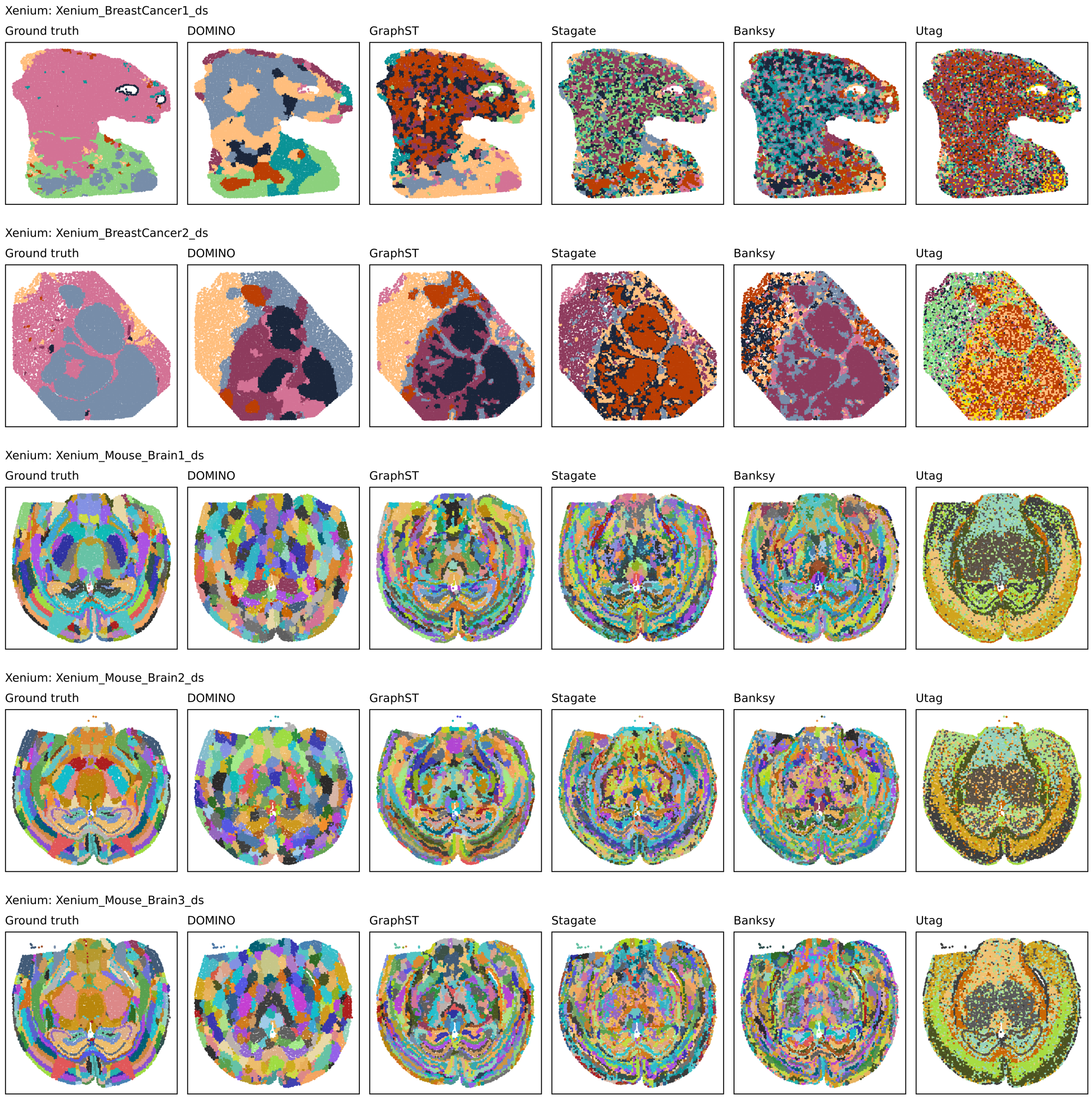


**Supplementary Figure 3.** Visualisation of spatial domains identification in Xenium datasets (down-sampled) from different platforms, with ground truth annotations shown alongside results from DOMINO, GraphST, STAGATE, Banksy, and UTAG.

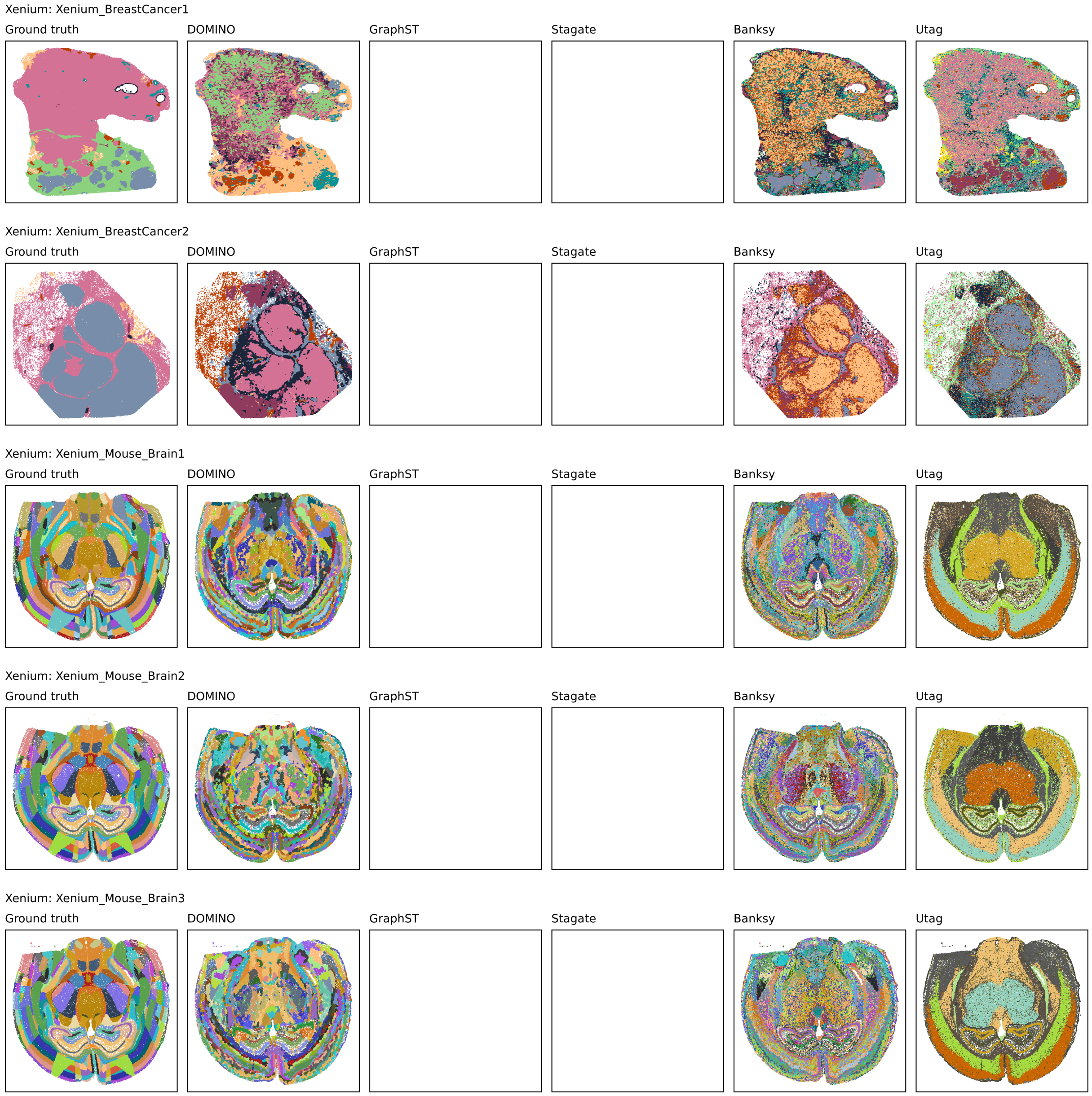


**Supplementary Figure 4.** Visualisation of spatial domains identification in Xenium datasets from different platforms, with ground truth annotations shown alongside results from DOMINO, GraphST, STAGATE, Banksy, and UTAG. Visualisation with a blank figure indicates that the method failed to process the ST data.

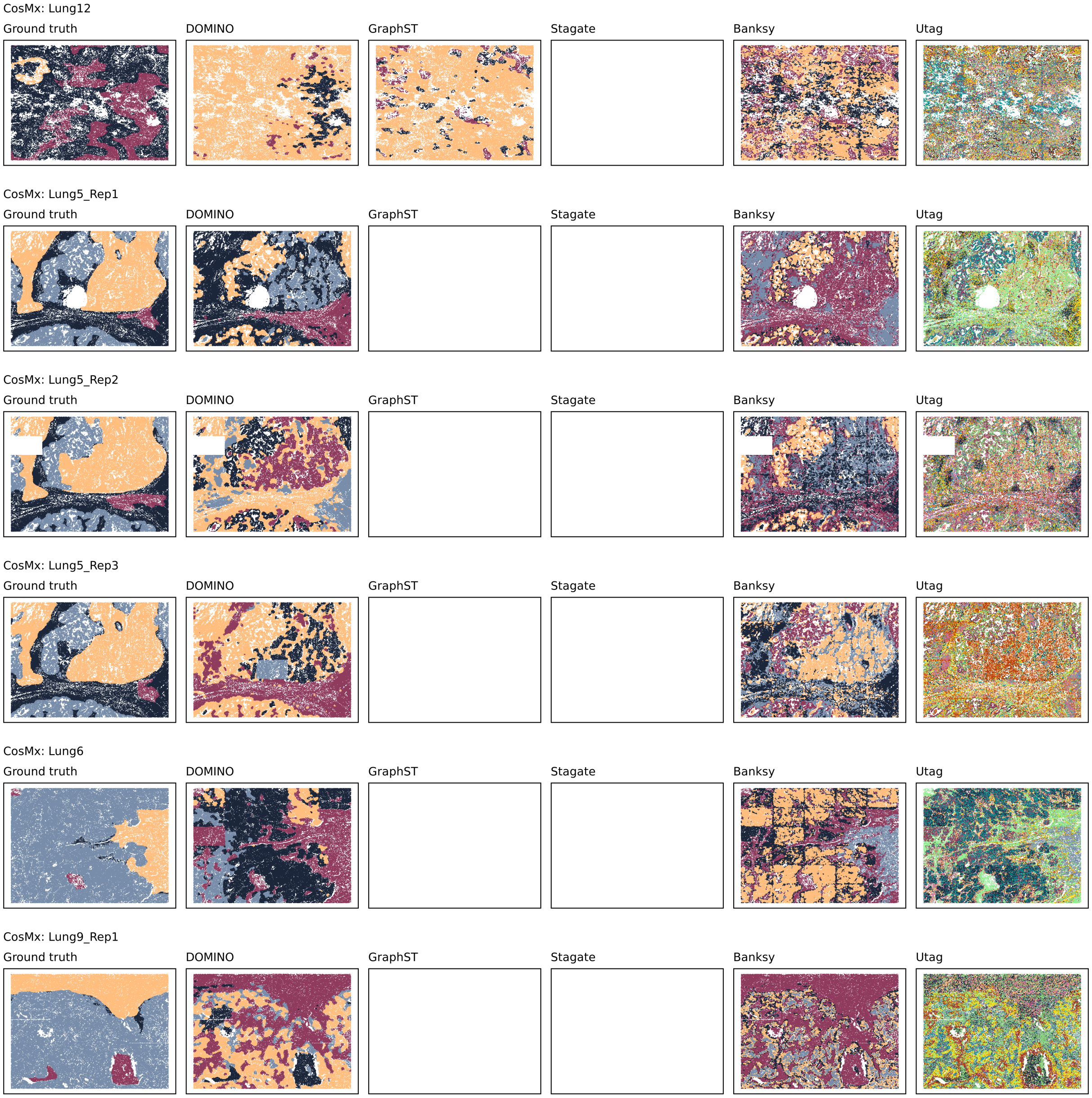


**Supplementary Figure 5.** Visualisation of spatial domains identification in CosMx datasets from different platforms, with ground truth annotations shown alongside results from DOMINO, GraphST, STAGATE, Banksy, and UTAG. Visualisation with a blank figure indicates that the method failed to process the ST data.

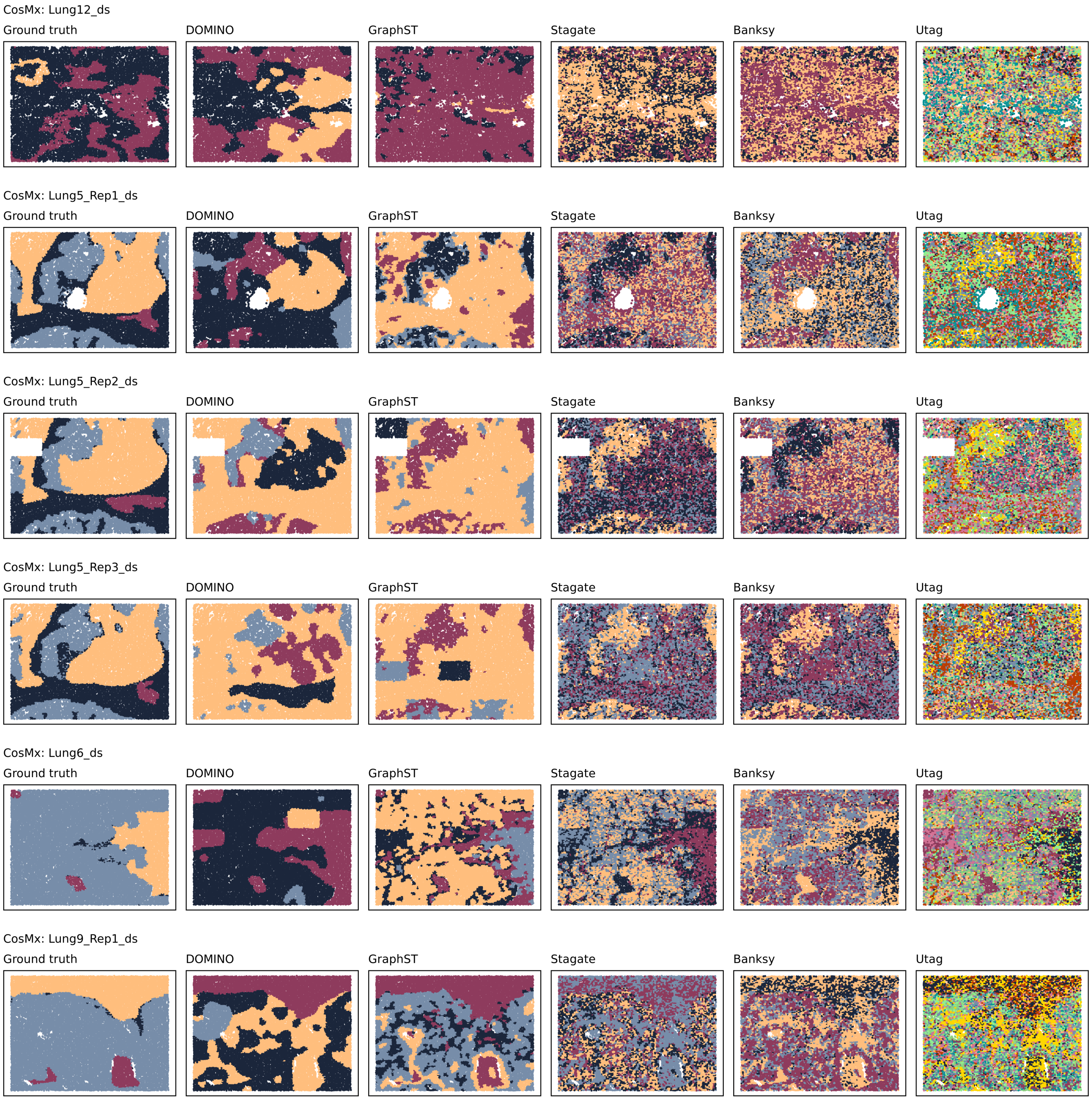


**Supplementary Figure 6.** Visualisation of spatial domains identification in CosMx datasets (down-sampled) from different platforms, with ground truth annotations shown alongside results from DOMINO, GraphST, STAGATE, Banksy, and UTAG.

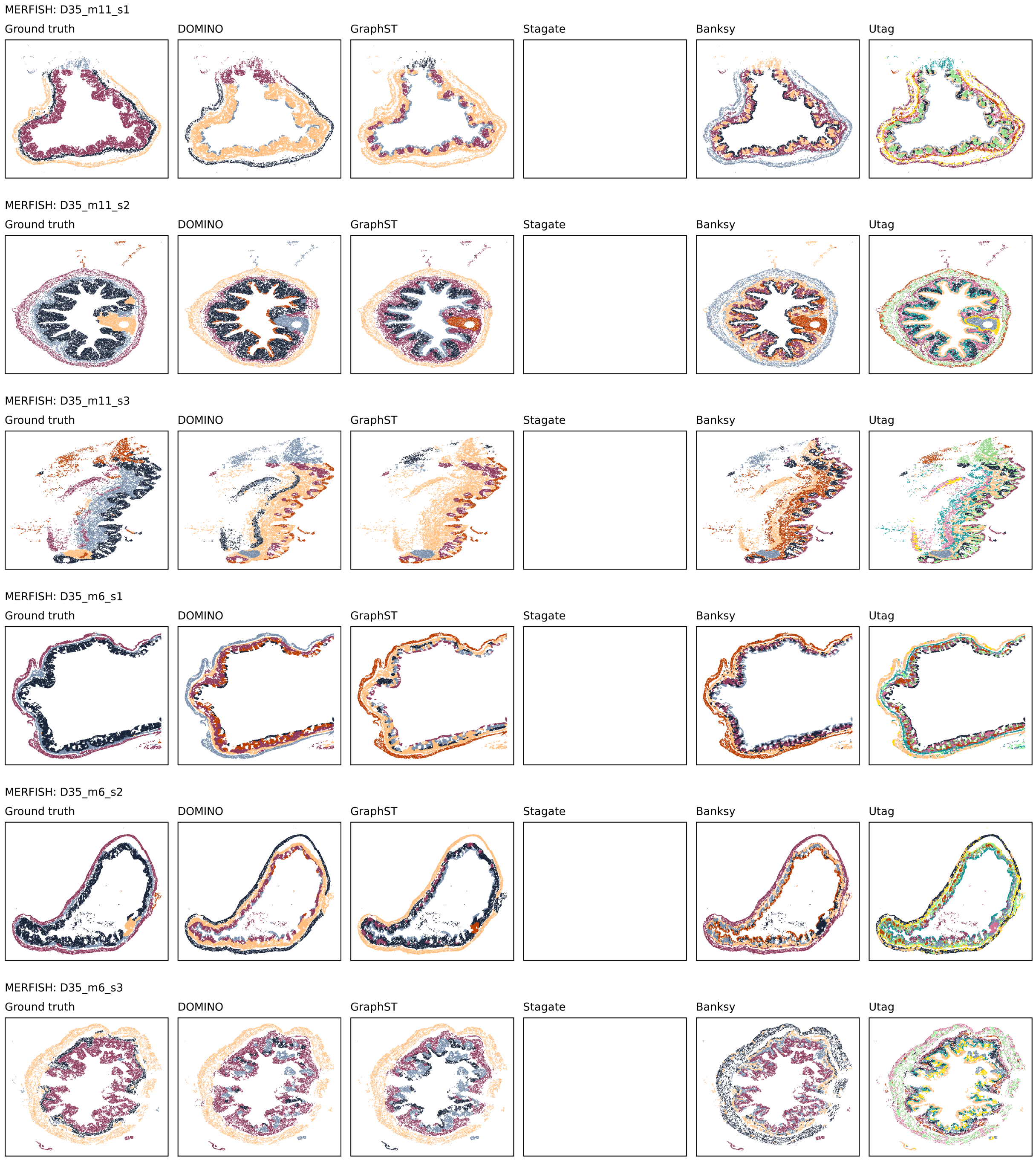


**Supplementary Figure 7.** Visualisation of spatial domains identification in MERFISH datasets from different platforms, with ground truth annotations shown alongside results from DOMINO, GraphST, STAGATE, Banksy, and UTAG. Visualisation with a blank figure indicates that the method failed to process the ST data.

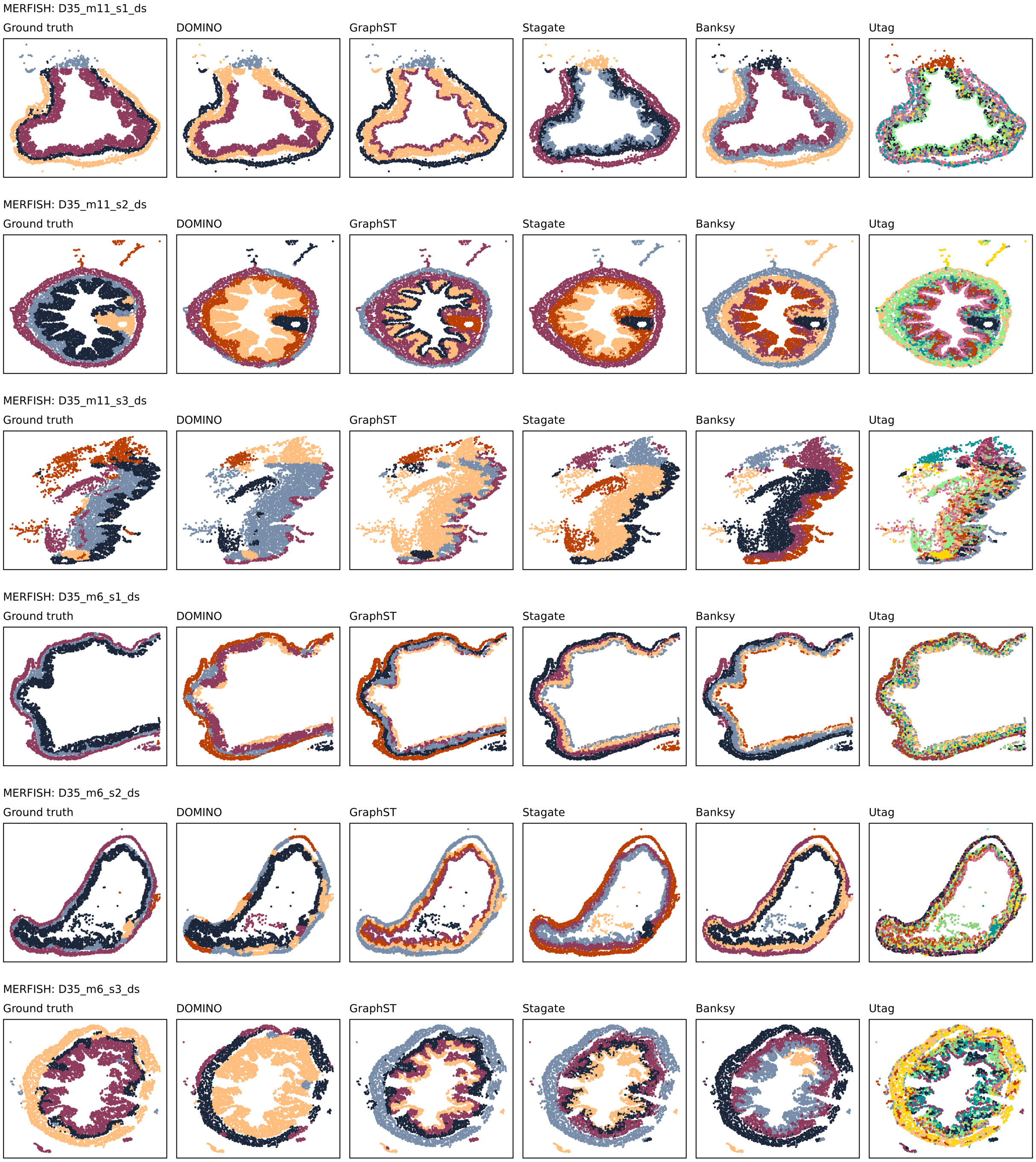


**Supplementary Figure 8.** Visualisation of spatial domains identification in MERFISH datasets (down-sampled) from different platforms, with ground truth annotations shown alongside results from DOMINO, GraphST, STAGATE, Banksy, and UTAG.

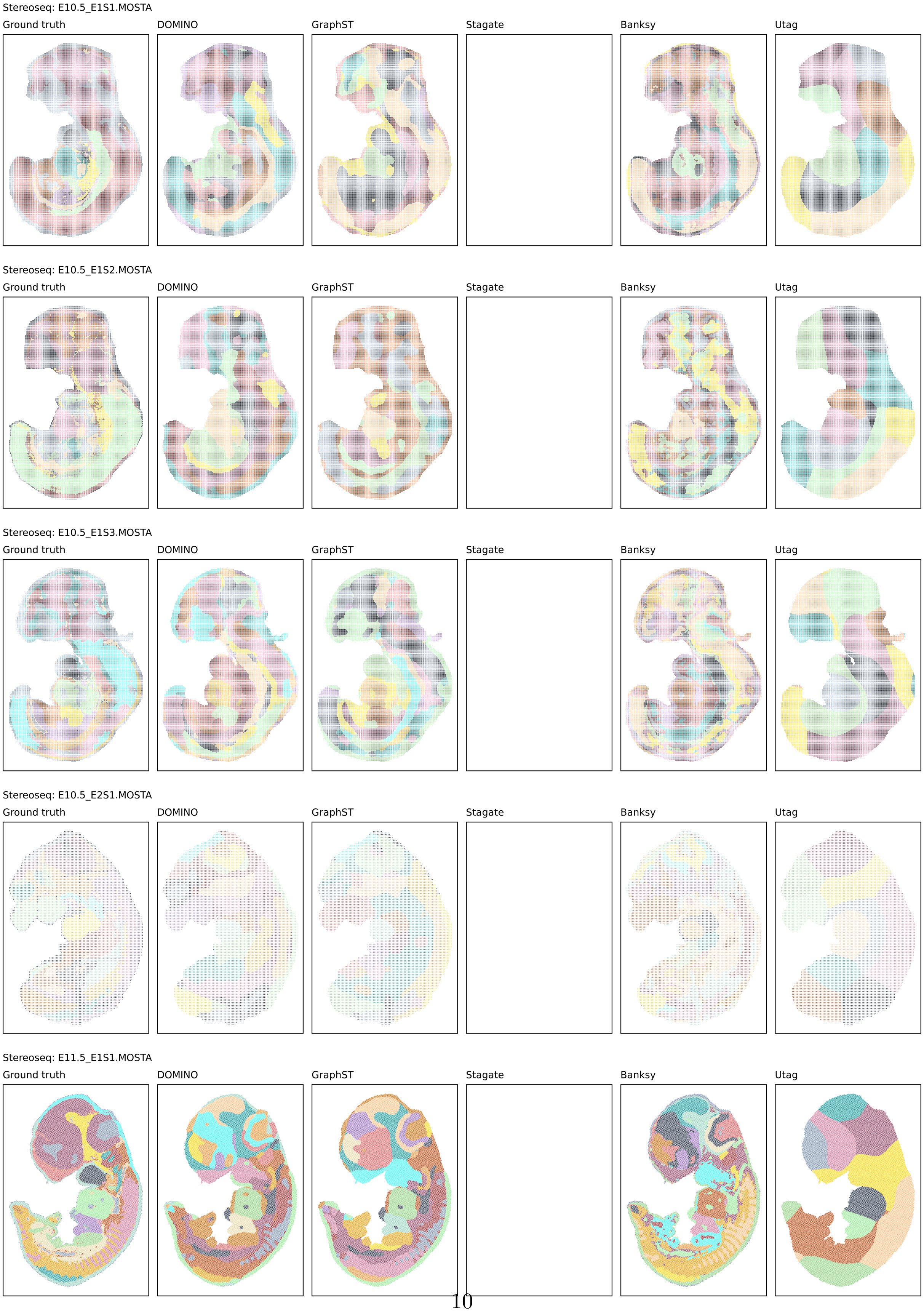




**Supplementary Figure 9.** Visualisation of spatial domains identification in Stereoseq datasets from different platforms, with ground truth annotations shown alongside results from DOMINO, GraphST, STAGATE, Banksy, and UTAG. Visualisation with a blank figure indicates that the method failed to process the ST data.

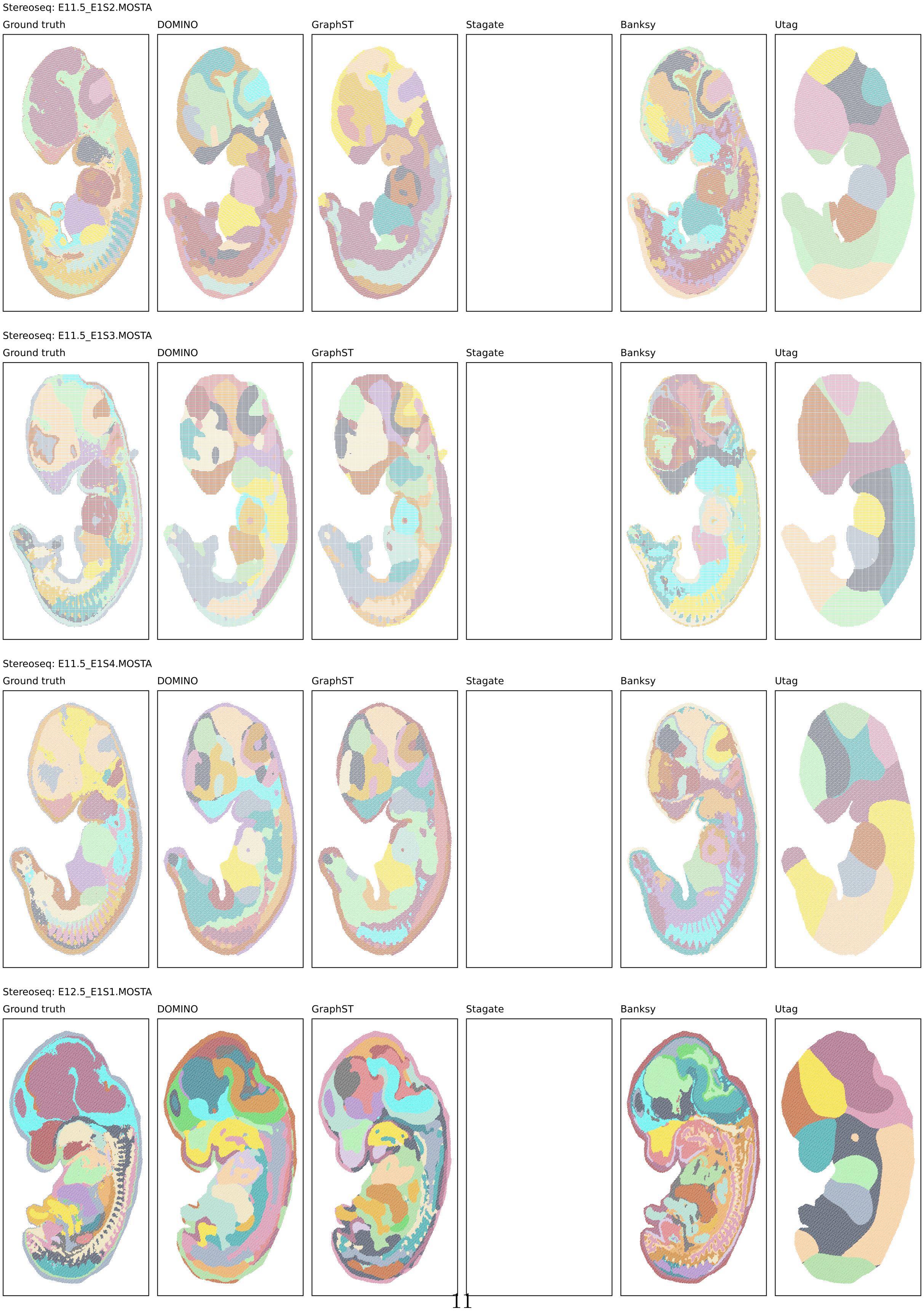




**Supplementary Figure 10.** Visualisation of spatial domains identification in Stereoseq datasets from different platforms, with ground truth annotations shown alongside results from DOMINO, GraphST, STAGATE, Banksy, and UTAG. Visualisation with a blank figure indicates that the method failed to process the ST data.

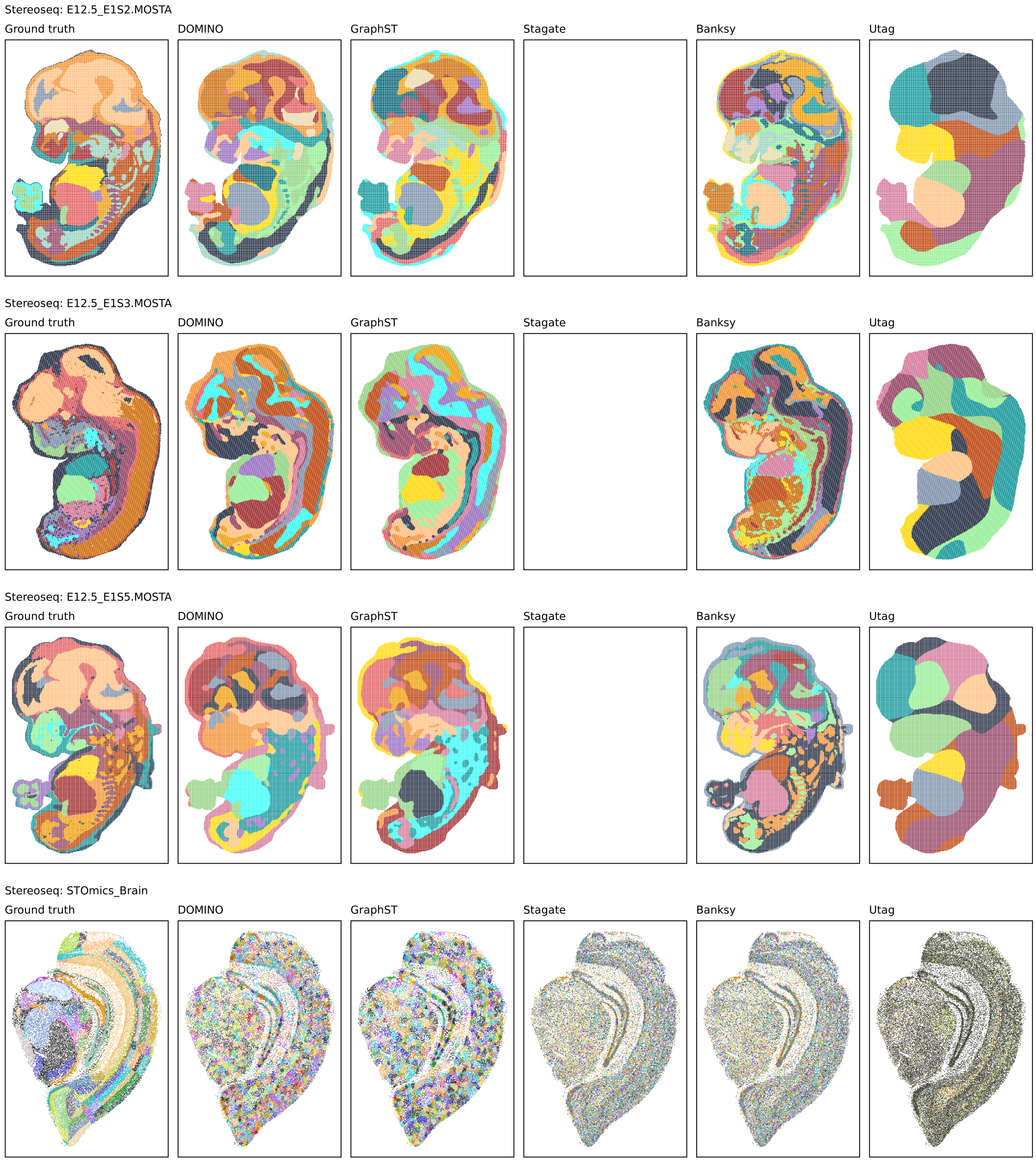


**Supplementary Figure 11.** Visualisation of spatial domains identification in Stereoseq datasets from different platforms, with ground truth annotations shown alongside results from DOMINO, GraphST, STAGATE, Banksy, and UTAG. Visualisation with a blank figure indicates that the method failed to process the ST data.

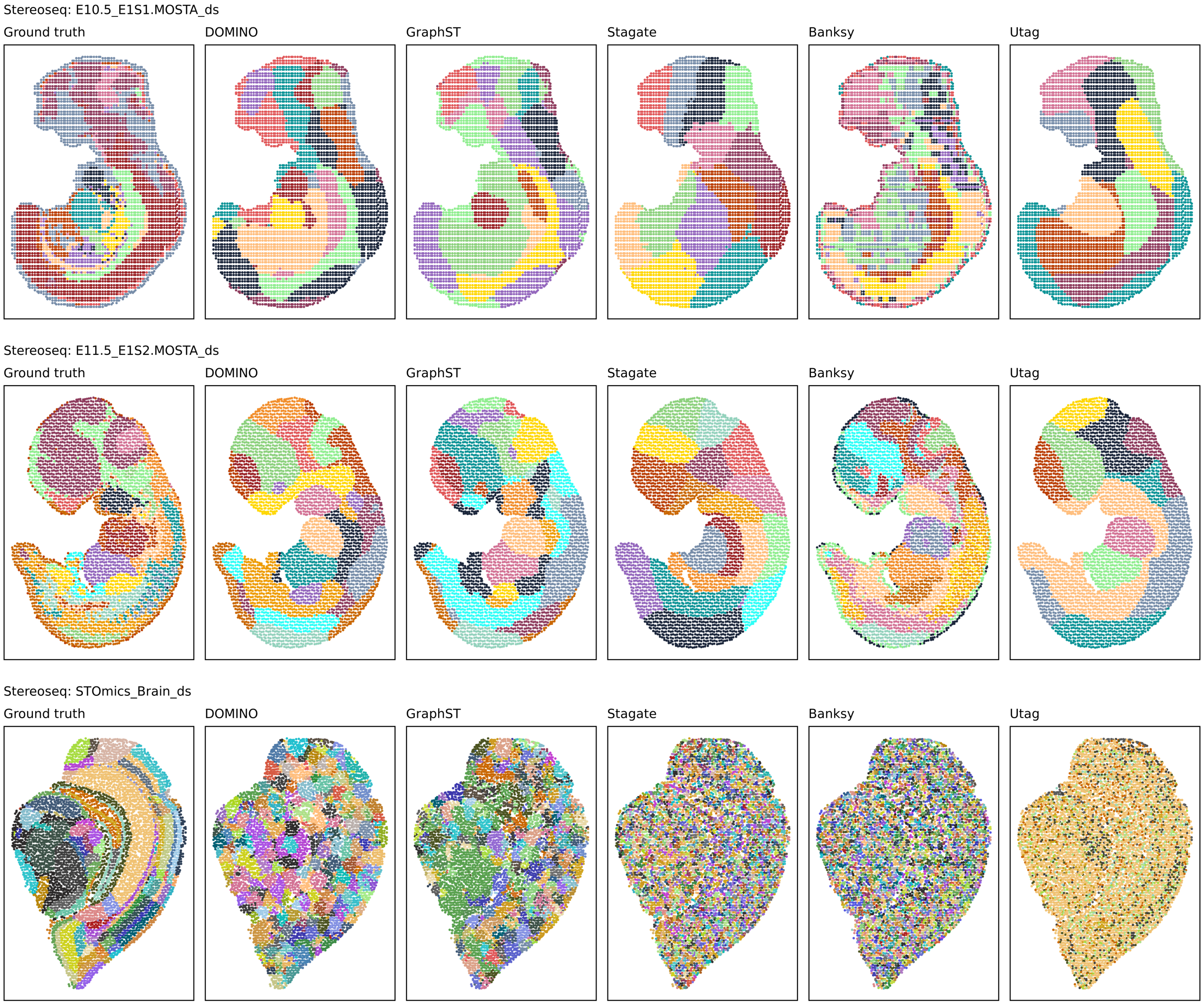


**Supplementary Figure 12.** Visualisation of spatial domains identification in Stereoseq datasets (down-sampled) from different platforms, with ground truth annotations shown alongside results from DOMINO, GraphST, STAGATE, Banksy, and UTAG.

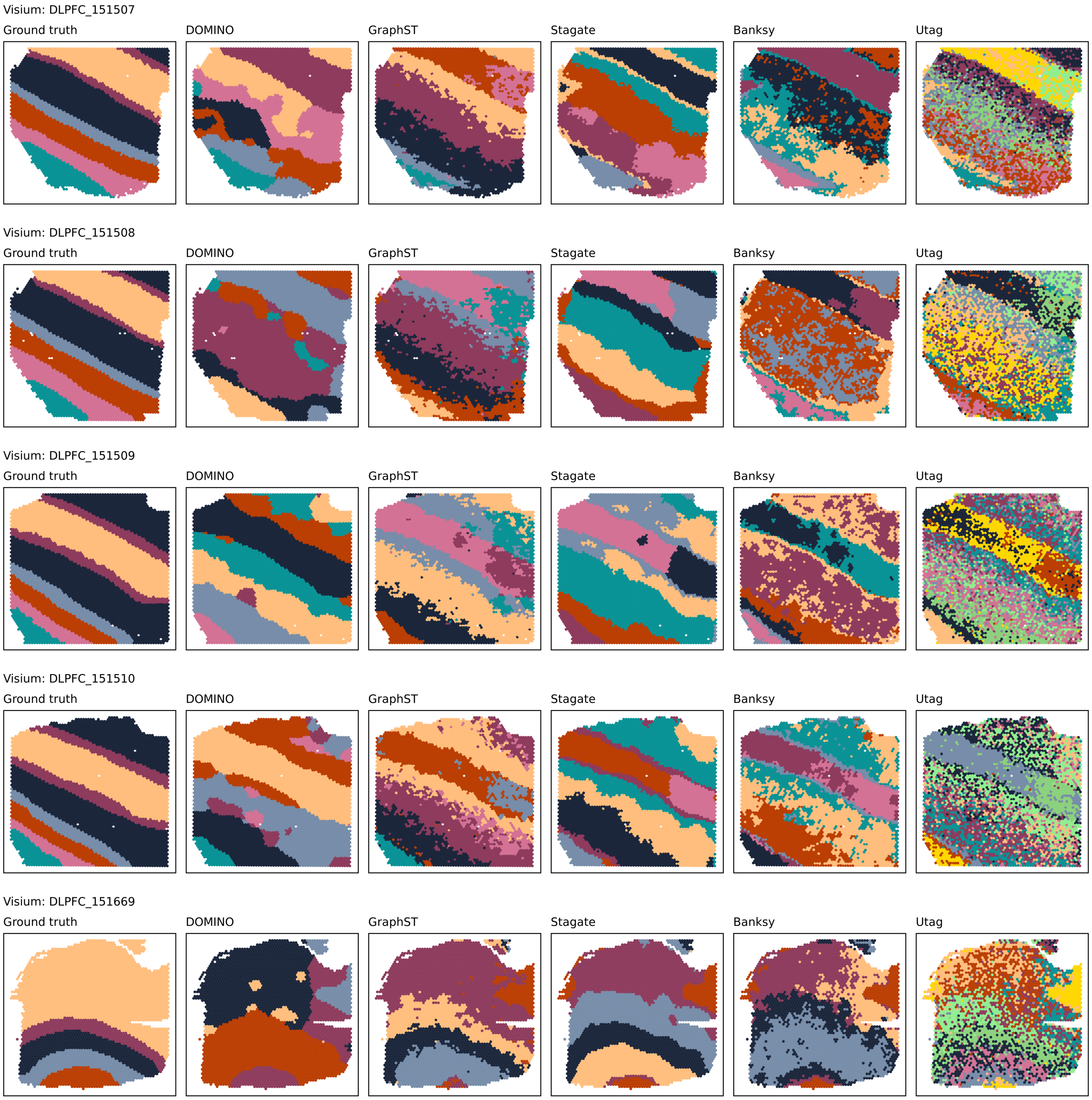


**Supplementary Figure 13.** Visualisation of spatial domains identification in Visium datasets from different platforms, with ground truth annotations shown alongside results from DOMINO, GraphST, STAGATE, Banksy, and UTAG.

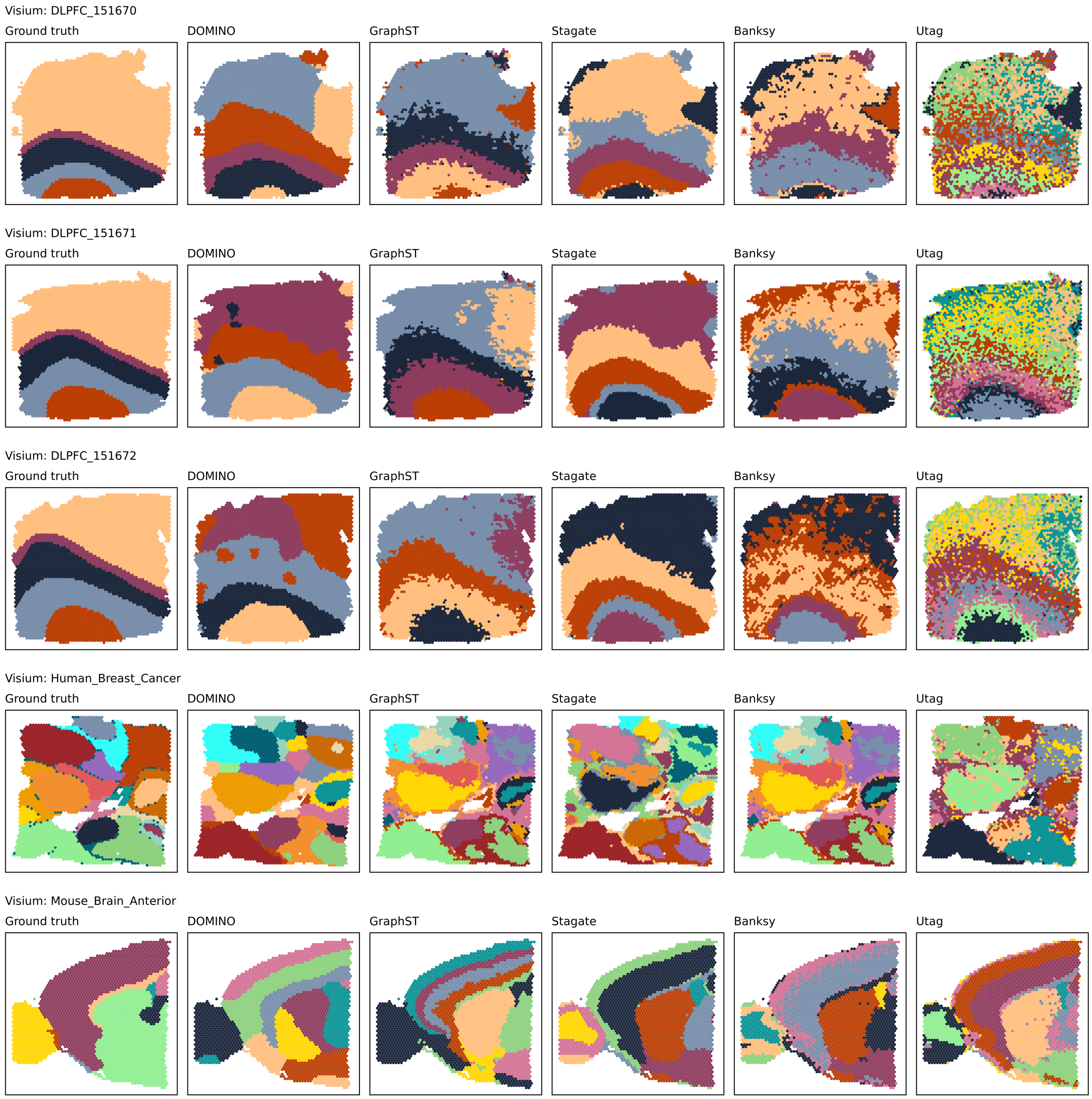


**Supplementary Figure 14.** Visualisation of spatial domains identification in Visium datasets from different platforms, with ground truth annotations shown alongside results from DOMINO, GraphST, STAGATE, Banksy, and UTAG.

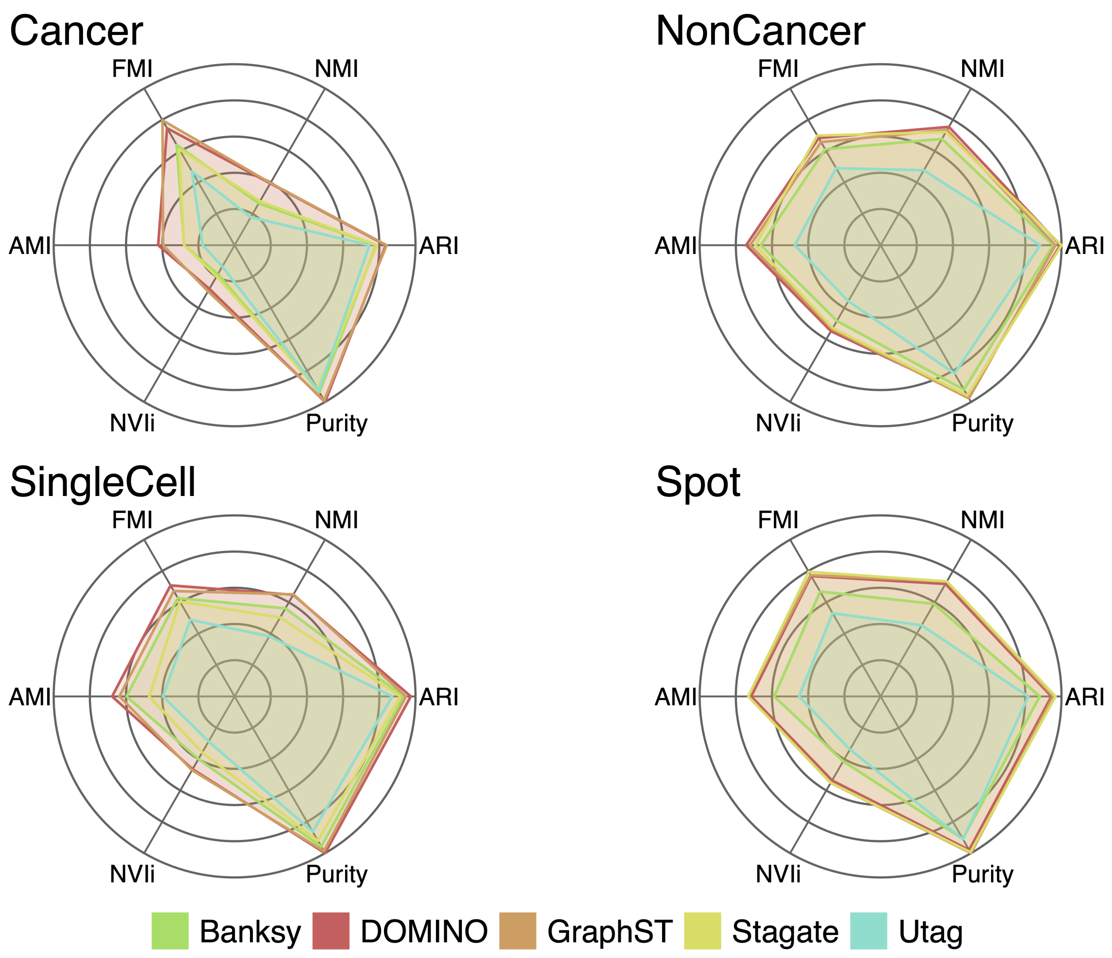


**Supplementary Figure 15.** Radar plots summarising six clustering accuracy metrics across five platforms (CosMx, MERFISH, Visium, Xenium, Stereo-seq) in different groups of datasets, including adjusted Rand index (ARI), normalised mutual information (NMI), adjusted mutual information (AMI), Fowlkes–Mallows index (FMI), inverted normalised variation of information (NVIi), and Purity.

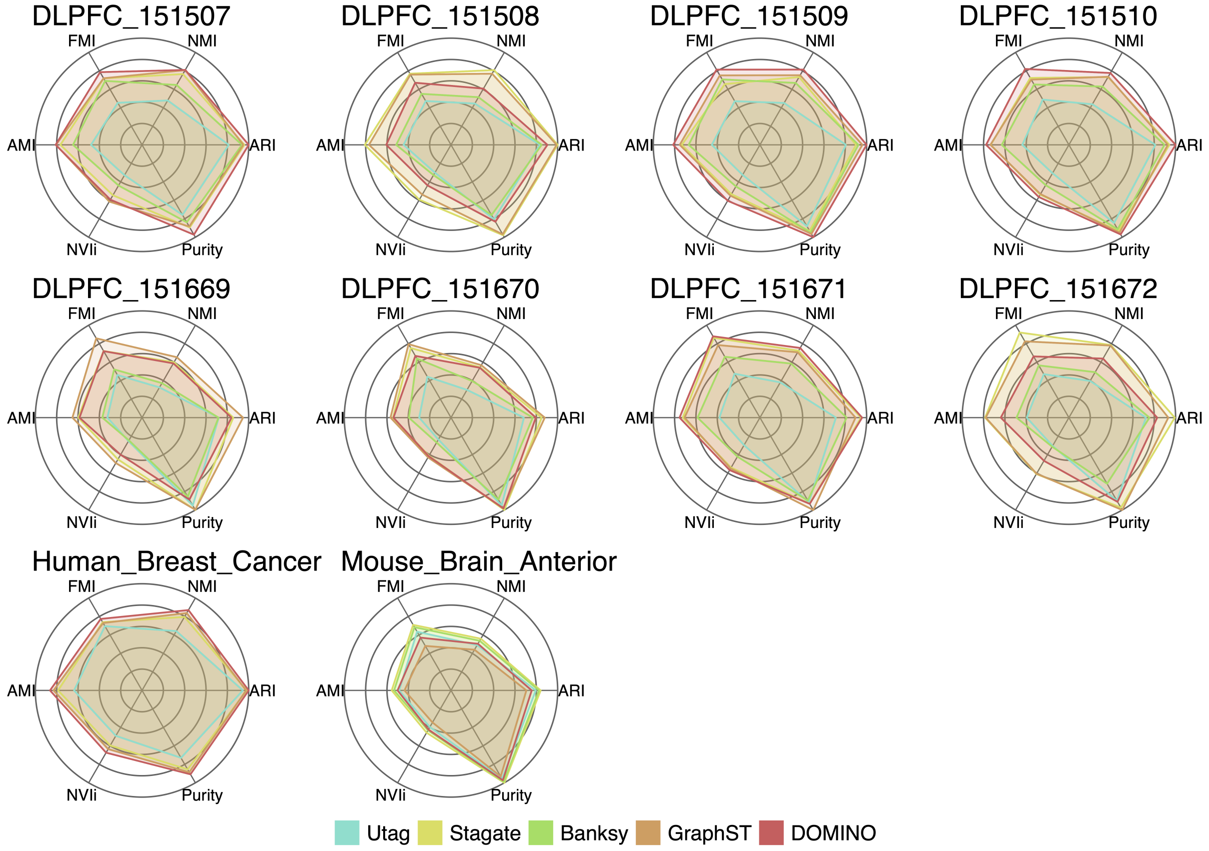


**Supplementary Figure 16.** Radar plots summarising six clustering accuracy metrics across five platforms (CosMx, MERFISH, Visium, Xenium, Stereo-seq) in Visium datasets, including adjusted Rand index (ARI), normalised mutual information (NMI), adjusted mutual information (AMI), Fowlkes–Mallows index (FMI), inverted normalised variation of information (NVIi), and Purity.

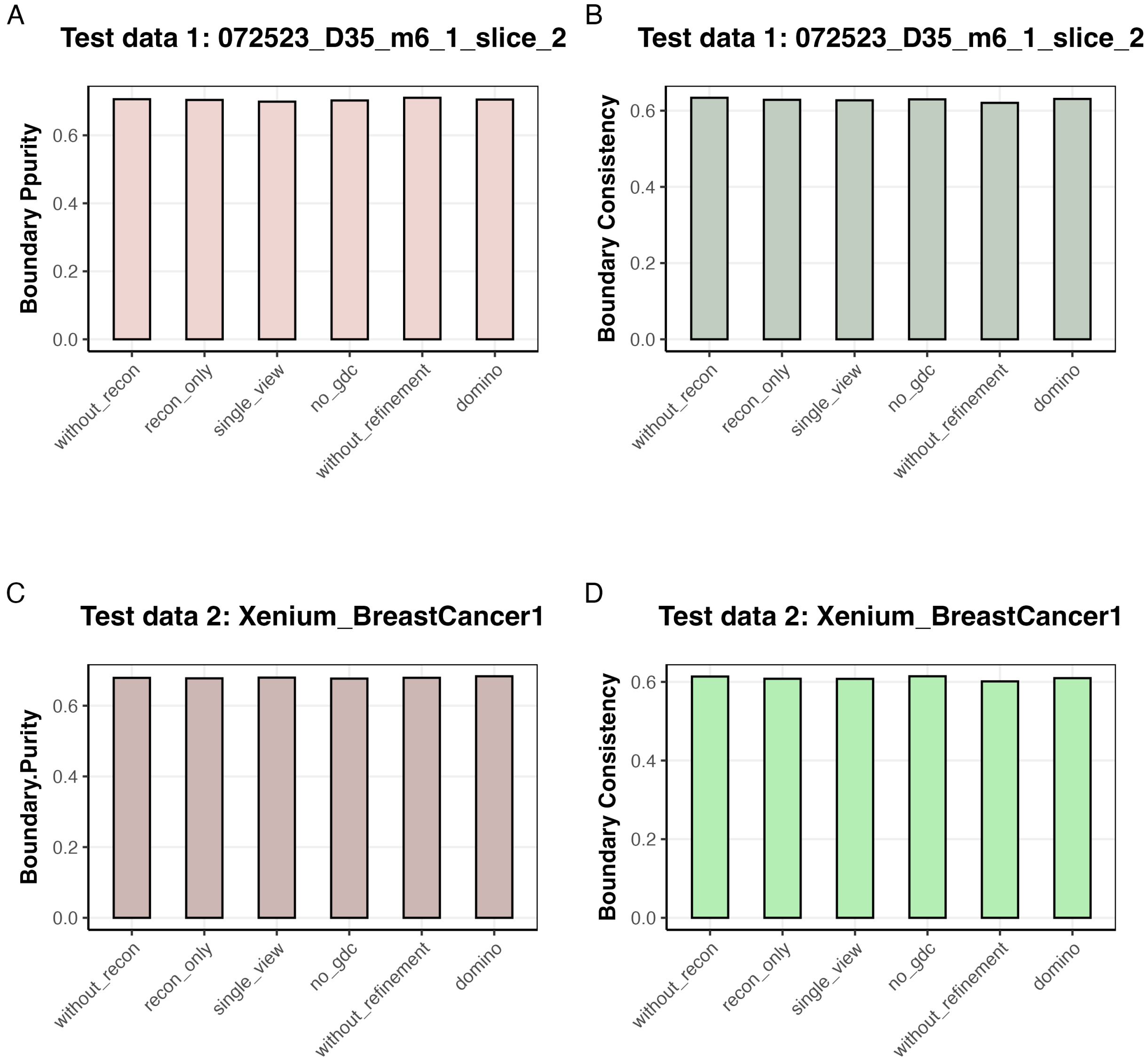


**Supplementary Figure 17.** Boundary metric comparison across DOMINO ablation variants. Boundary purity (BP; higher is better) and boundary consistency (BC; higher is better) were evaluated across DOMINO ablation settings on two test datasets. **a,b**, BP and BC for test data 1 (*072523_D35_m6_1_slice_2*). **c,d**, BP and BC for test data 2 (*Xenium_BreastCancer1*). The ablation settings included: *without_recon*, model trained without feature reconstruction loss; *recon_only*, reconstruction-only baseline without graph contrastive learning; *single_view*, single-view graph contrastive model; *no_gdc*, model in which the graph diffusion view was replaced by an alternative augmentation strategy; *without_refinement*, full DOMINO model without the post-clustering refinement step; and *domino*, the complete DOMINO model. Across both datasets, the full DOMINO model showed the strongest overall boundary performance, whereas the refined and unrefined variants yielded highly similar BP and BC values, indicating that the improved boundary quality is primarily attributable to the learned domain structure rather than post hoc label refinement.

Benchmarks for boundary fidelity and fine-grained structure recovery

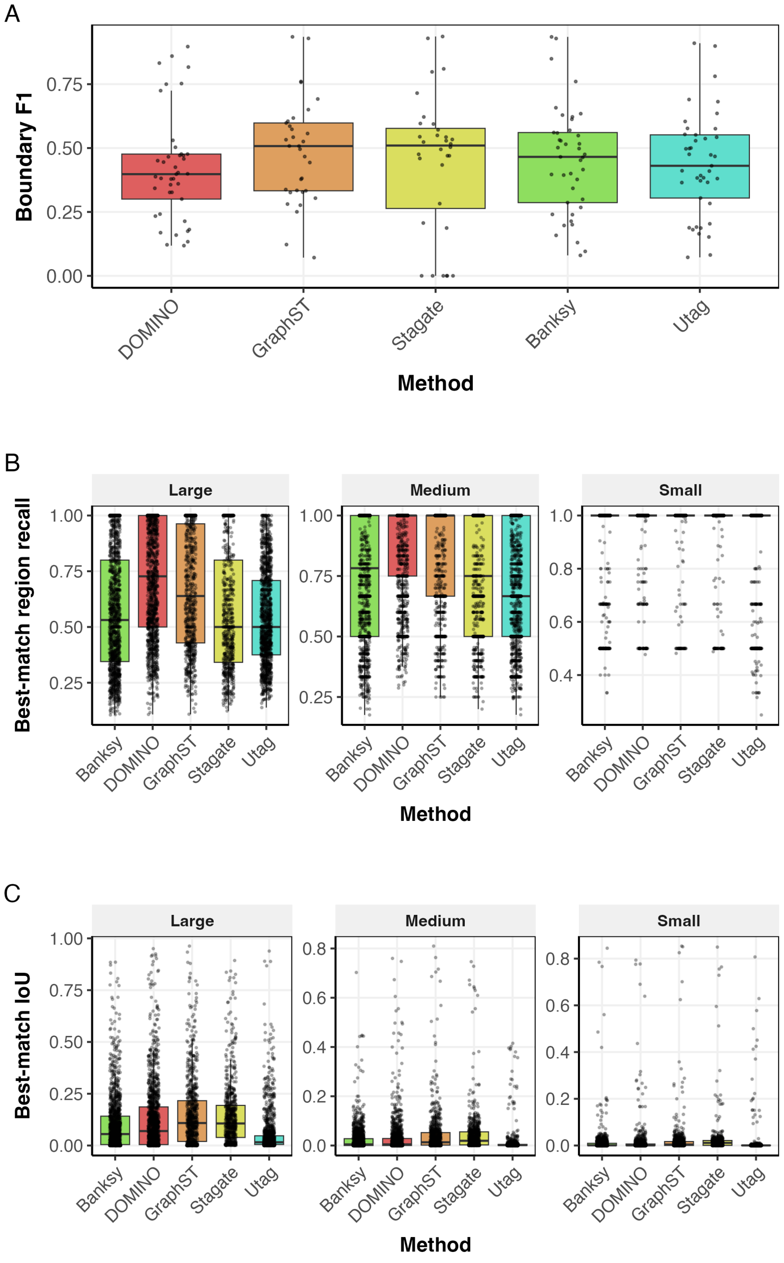


**Supplementary Figure 18.** Benchmarking boundary fidelity and fine-grained structure recovery across methods. **a**, Boundary-level agreement between predicted and annotated domain interfaces, measured by boundary F1 across benchmark datasets. **b**, Recovery of annotated regions measured by best-match recall, stratified by region size (*large*, *medium* and *small*). For each annotated region, the predicted region with the highest overlap was identified, and recall was calculated relative to the annotated region. **c**, Recovery of annotated regions measured by best-match intersection over union (IoU), also stratified by region size. Higher values indicate better recovery of annotated structures. DOMINO achieved competitive boundary F1 scores and maintained strong recovery of small and medium-sized regions relative to other methods, indicating that its improved spatial coherence is not explained by simply merging domains into larger blocks, but is accompanied by preservation of boundary fidelity and fine-scale tissue structures.

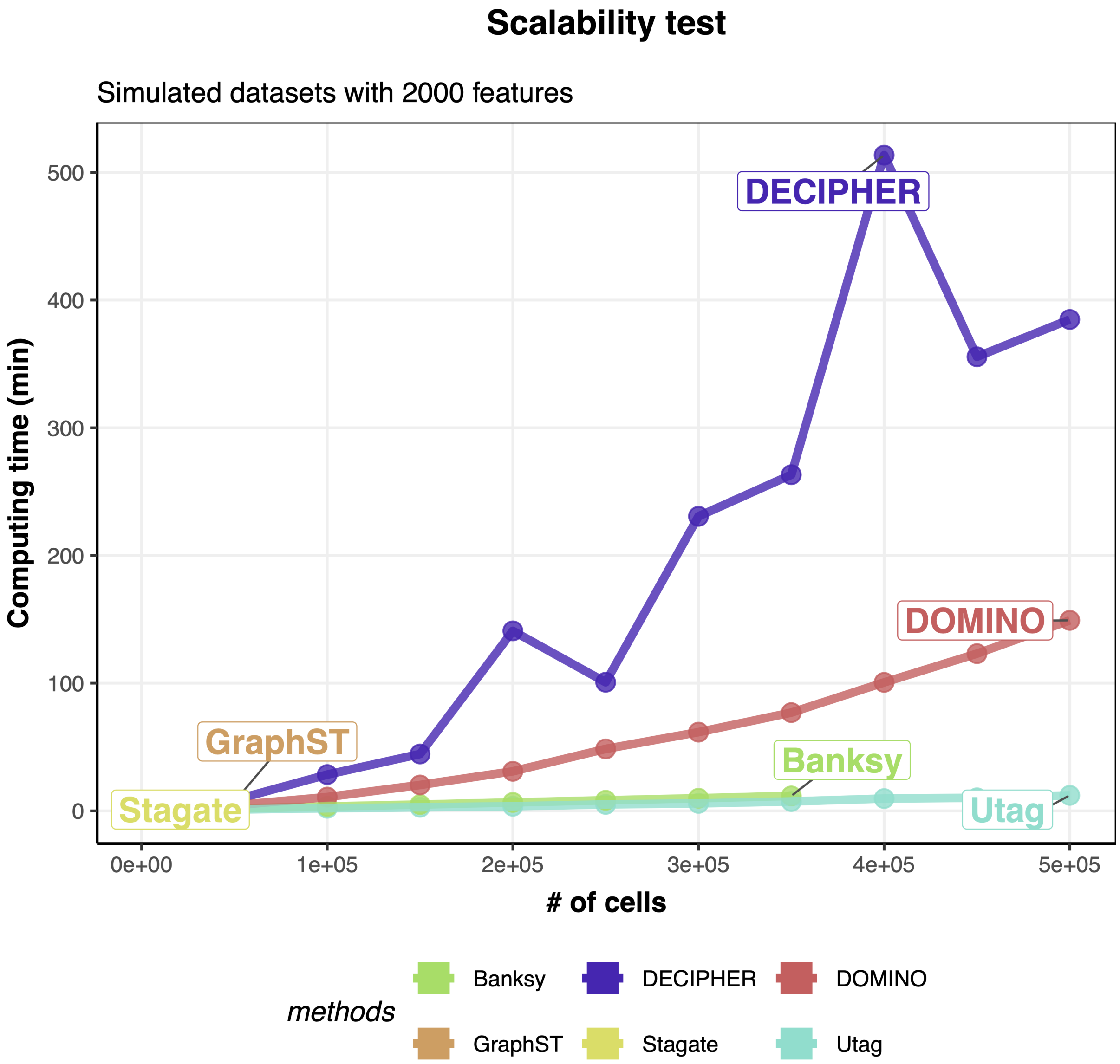


**Supplementary Figure 19.** Scalability benchmarking across simulated spatial transcriptomics datasets. Computing time was evaluated for DOMINO and five comparison methods on simulated spatial transcriptomics datasets ranging from 1,000 to 500,000 cells, each with 2,000 features. All methods were benchmarked under a standardised computing environment comprising a 14-core CPU, 100 GB RAM and an A800 GPU with 80 GB VRAM. DOMINO showed stable scalability across increasing dataset sizes, whereas Banksy encountered a memory outage at 400,000 cells, and both GraphST and Stagate failed beyond 100,000 cells. DECIPHER, a method designed for large-scale spatial transcriptomics analysis, was included as an additional comparator.

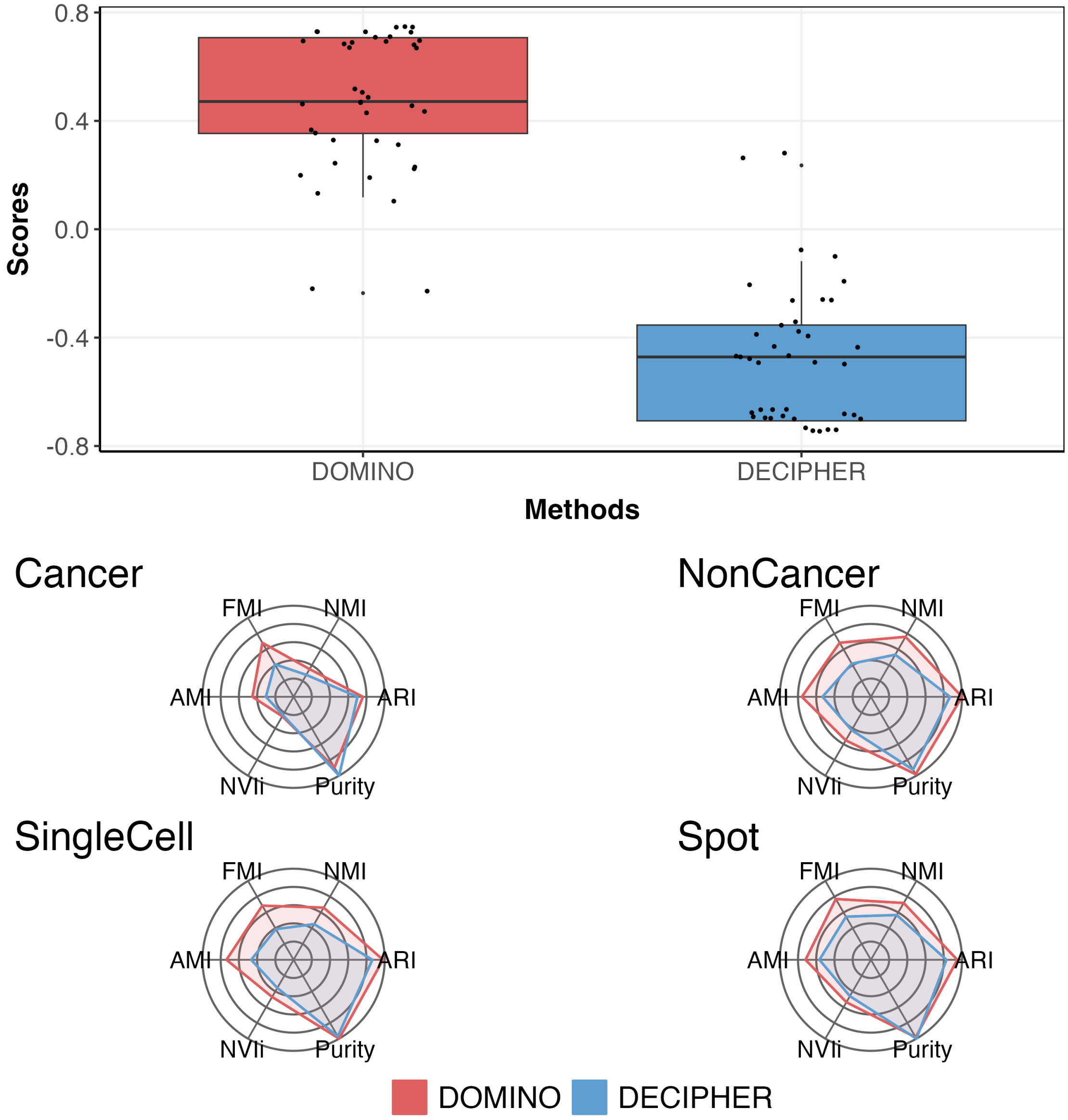


**Supplementary Figure 20.** Accuracy comparison between DOMINO and DECIPHER across benchmark datasets. **a**, Distribution of overall performance scores for DOMINO and DECIPHER across benchmark datasets. **b**, Radar plots comparing mean performance across six evaluation metrics (ARI, NMI, FMI, AMI, NVIi and purity) in cancer and non-cancer datasets. **c**, Radar plots comparing mean performance across the same six metrics in single-cell and spot-based spatial transcriptomics datasets. DOMINO showed higher overall performance than DECIPHER across benchmark datasets and dataset categories.

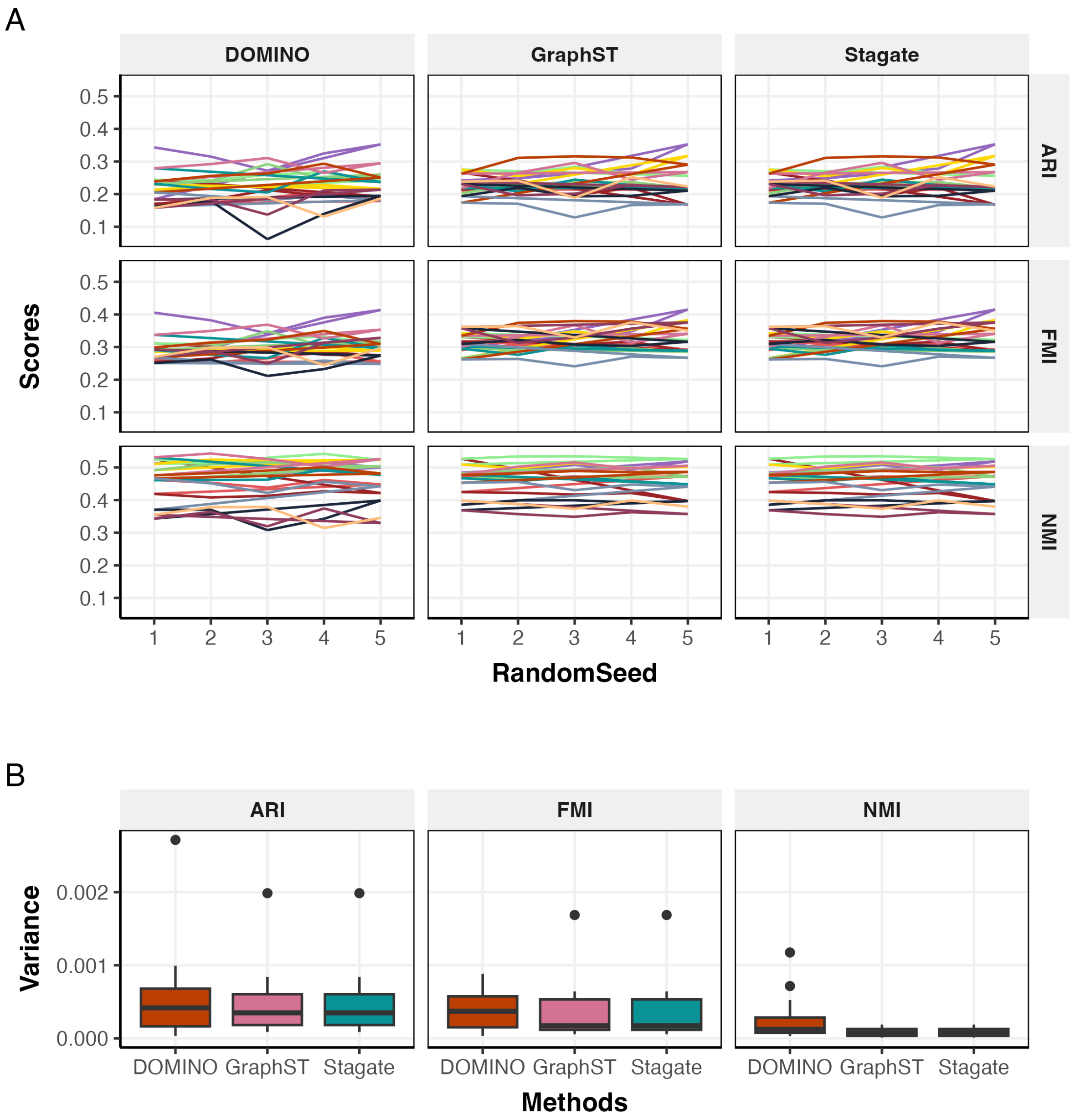


**Supplementary Figure 21.** Stability of DOMINO across repeated runs with different random seeds.**a**, ARI, FMI and NMI scores obtained across five independent runs with different random seeds for DOMINO, GraphST and Stagate. Each coloured line represents one benchmark dataset, showing the variation in performance across repeated runs. **b**, Variance of ARI, FMI and NMI across the five runs for each method. DOMINO showed low variance across repeated runs, broadly comparable to other graph-based methods, indicating that its performance is not unusually sensitive to random initialisation or stochastic sampling during training.

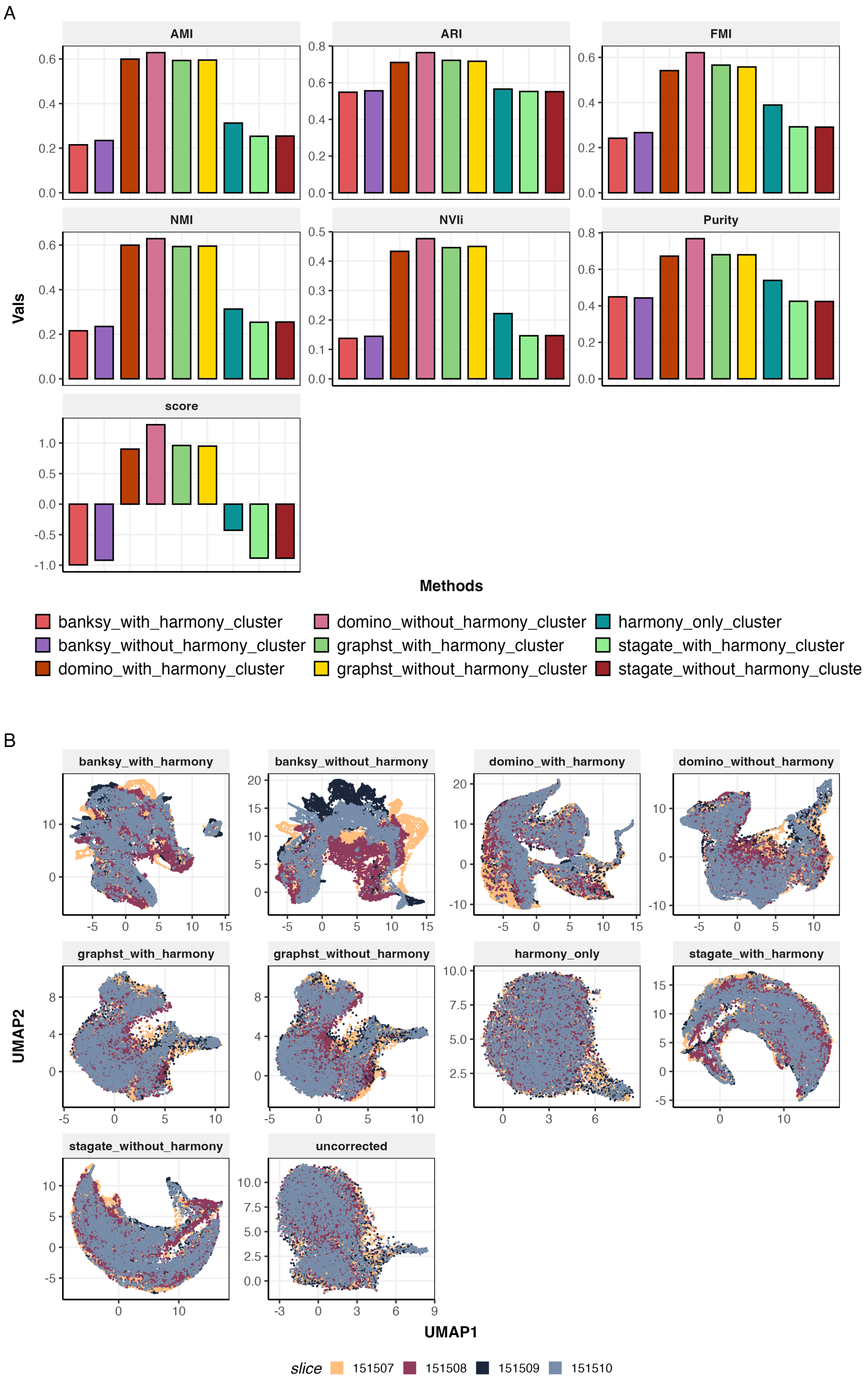


**Supplementary Figure 22.** Multi-sample domain identification in 4 DLPFC datasets. **a**, Performance comparison across methods and integration strategies, evaluated using AMI, ARI, FMI, NMI, NVIi, purity and the overall score. Methods were applied either directly, after Harmony-based batch correction, or using Harmony alone followed by clustering. **b**, UMAP visualisation of the corresponding embeddings coloured by sample identity, showing the degree of sample mixing under each strategy. In this dataset, where batch effects between samples were minimal, DOMINO without Harmony achieved the best overall domain identification performance, indicating that additional non-spatial batch correction was not required.

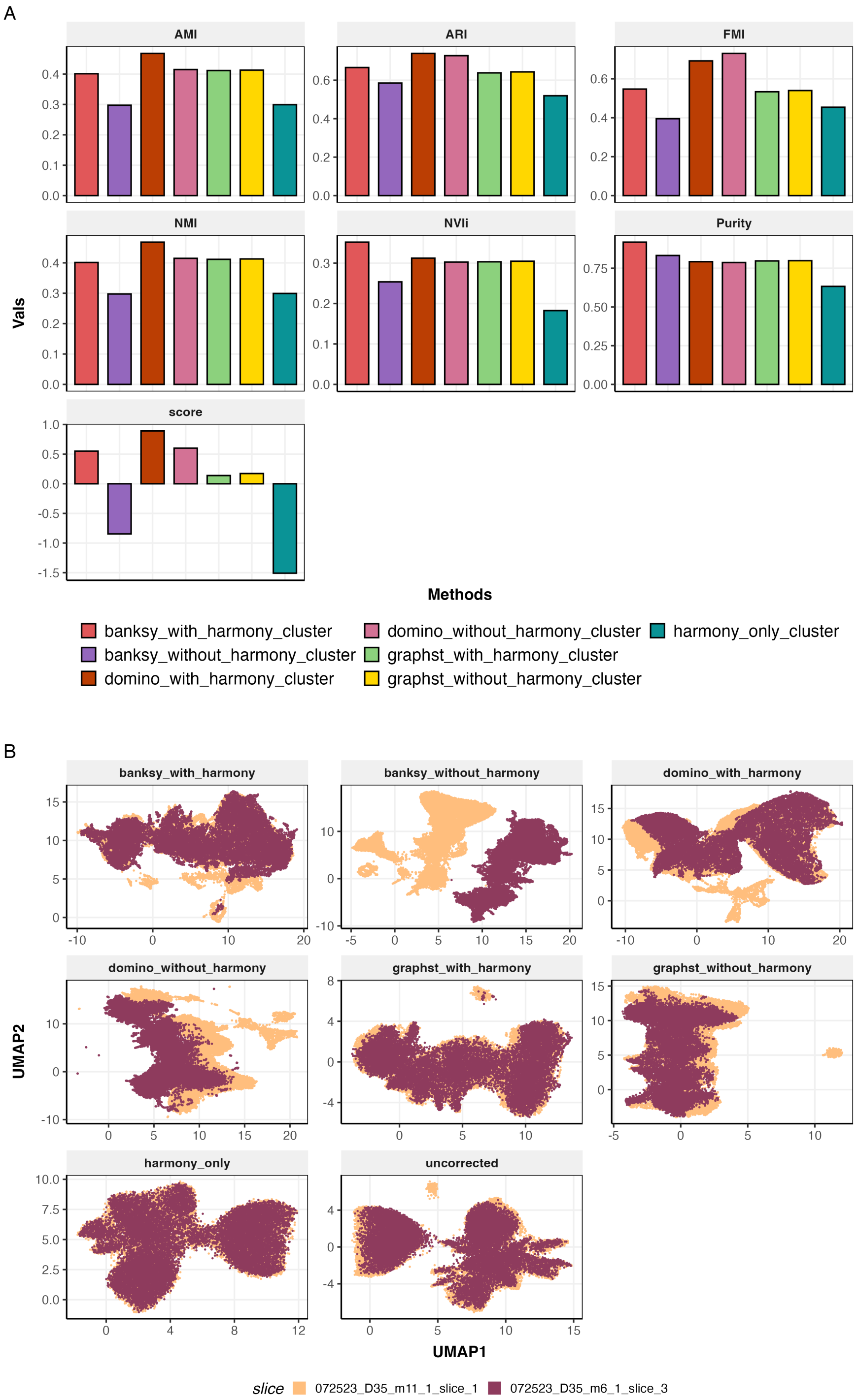


**Supplementary Figure 23.** Multi-sample domain identification in MERFISH mouse colon datasets with batch effects across two samples. **a**, Performance comparison across methods and integration strategies, evaluated using AMI, ARI, FMI, NMI, NVIi, purity and the overall score. Methods were applied with or without Harmony-based batch correction, alongside Harmony alone followed by clustering. **b**, UMAP visualisation of the corresponding embeddings coloured by sample identity. In this dataset, Harmony-corrected DOMINO embeddings followed by joint clustering achieved the best overall performance, indicating that DOMINO supports effective multi-sample integration when batch effects are present.

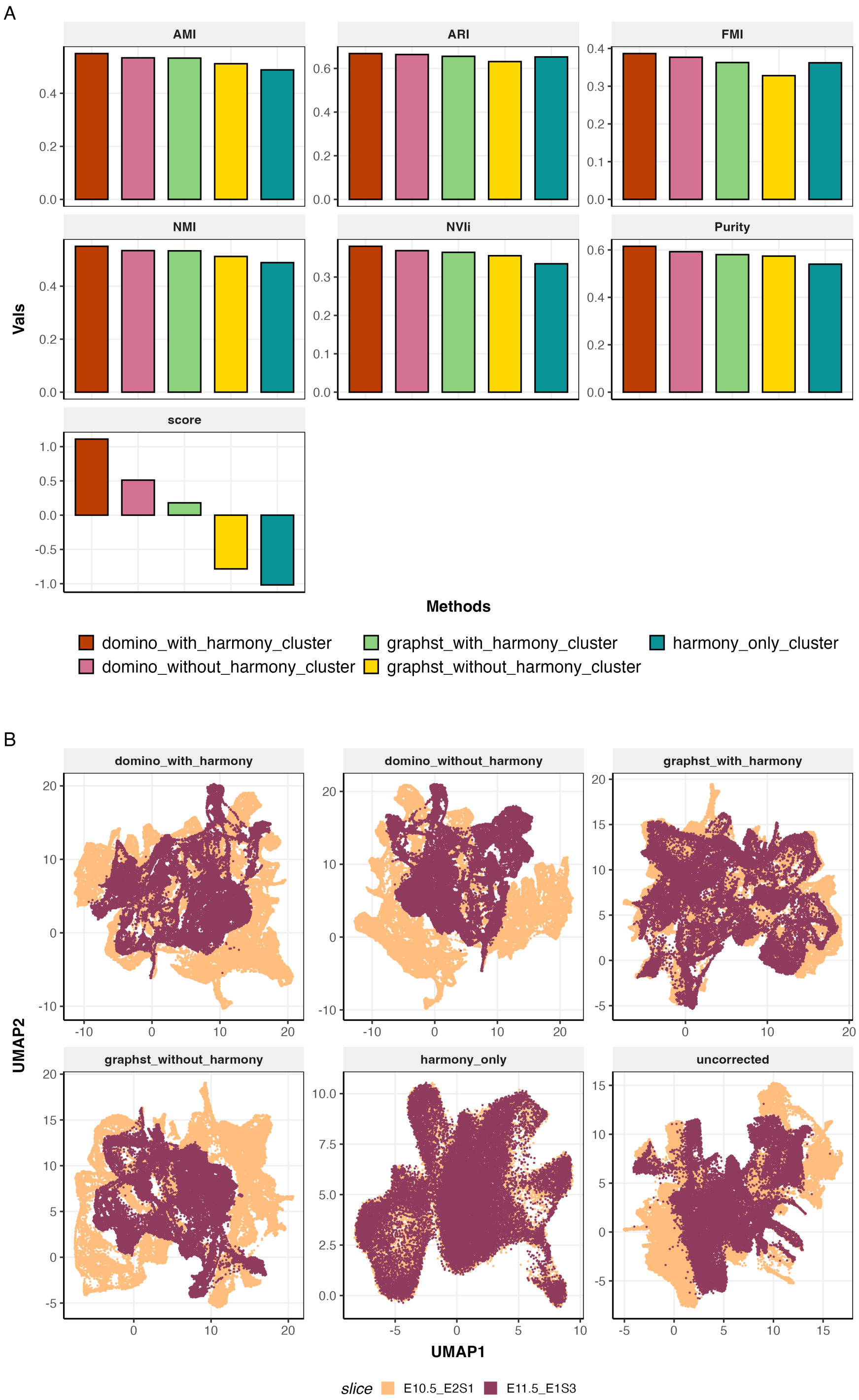


**Supplementary Figure 24.** Multi-sample domain identification in a Stereo-seq dataset with batch effects across two slices. **a**, Performance comparison across methods and integration strategies, evaluated using AMI, ARI, FMI, NMI, NVIi, purity and the overall score. Methods were applied with or without Harmony-based batch correction, alongside Harmony alone followed by clustering. **b**, UMAP visualisation of the corresponding embeddings coloured by slice identity. Harmony-corrected DOMINO embeddings yielded the highest overall performance, supporting the use of DOMINO for joint spatial domain identification across samples while accounting for batch effects.

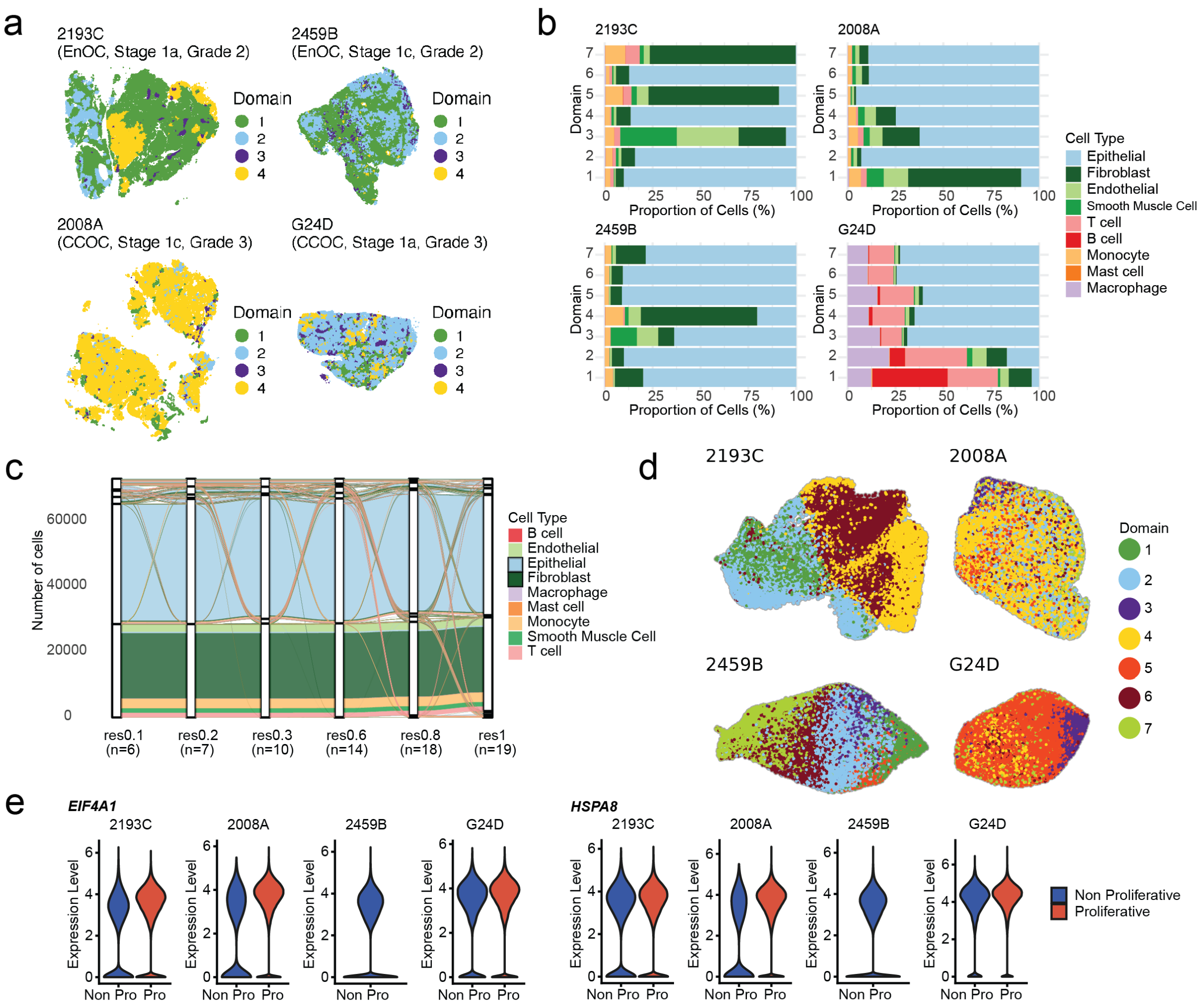


**Supplementary Figure 25. Additional analyses of spatial domains and clinical associations in EAOC.**(a) Spatial maps of four representative EAOC samples coloured by DOMINO domain assignments at a resolution of four domains per sample. (b) Proportion of cells from each cell type within each domain across all samples at a resolution of seven domains per sample. (c) Alluvial plot comparing BANKSY domain assignments at different resolution, illustrating that BANKSY does not further resolve tumour cells into distinct transcriptional states. (d) UMAP of tumour cells from each sample coloured by DOMINO domain assignment. (e) Violin plots showing log normalised expression of marker genes up-regulated in proliferative tumour domains.

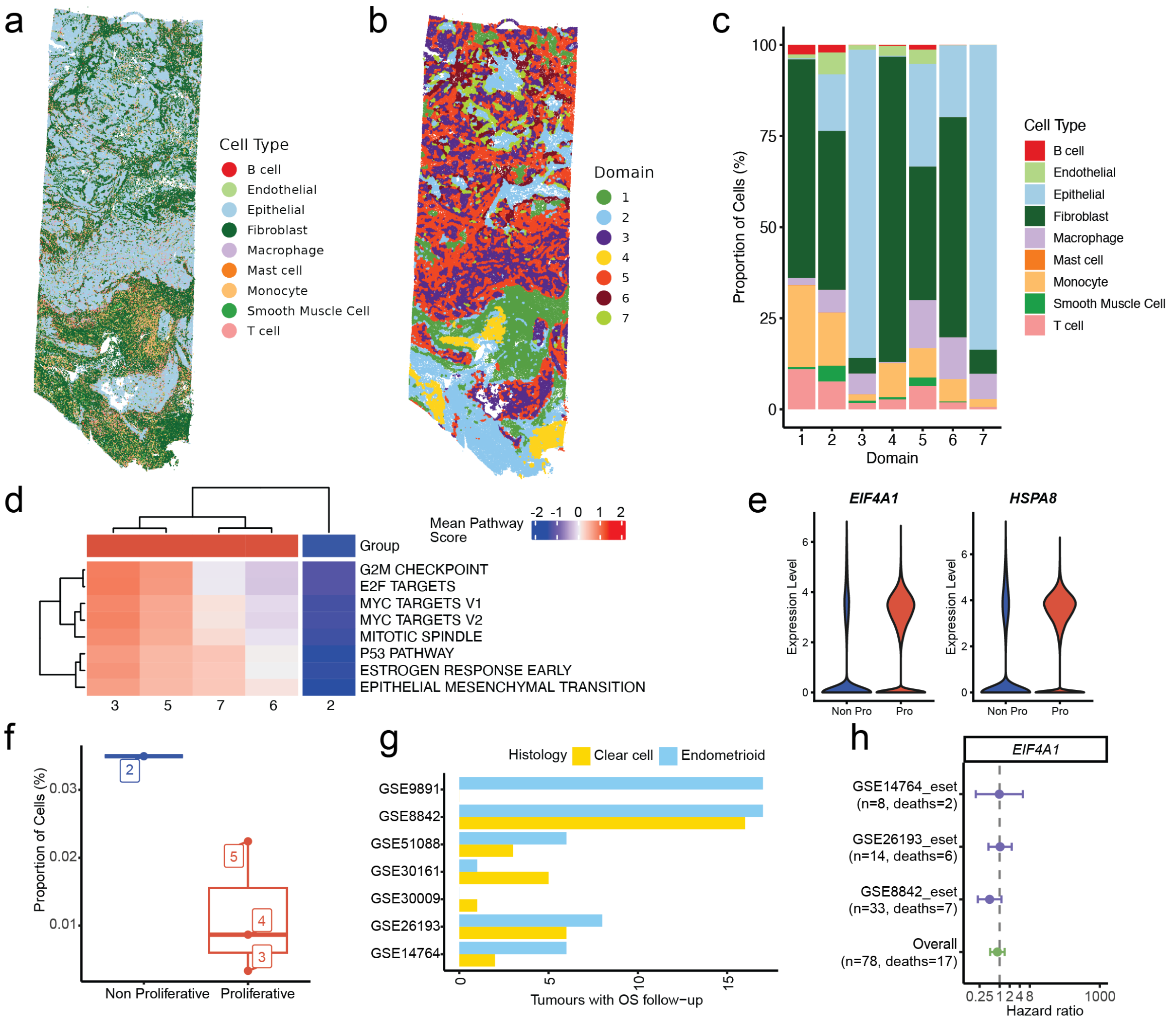


**Supplementary Figure 26. Validation of proliferative and non proliferative tumour states in an independent CCOC spatial transcriptomic dataset.** Spatial map of CCOC Xenium 5K data from Le et al.[?], colored by (a) cell type and (b) DOMINO domain assignment. (c) Proportion of cells from each cell type assigned to each of the seven DOMINO domains. (d) Hallmark pathway activity scores for proliferation associated pathways in epithelial cells from tumour enriched domains. (e) Violin plots showing increased expression of *EIF4A1* and *HSPA8* in epithelial cells from proliferative domains compared with the non proliferative domain. (f) Proportion of mast cells across proliferative and non proliferative tumour enriched domains. (g) Number of tumours from each ovarian cancer subtype with overall survival follow up available in publicly accessible datasets. (h) Forest plot showing hazard ratios and 95% confidence intervals from Cox proportional hazards analyses of *EIF4A1* expression across multiple ovarian cancer cohorts. No statistically significant association with overall survival was observed in any dataset.

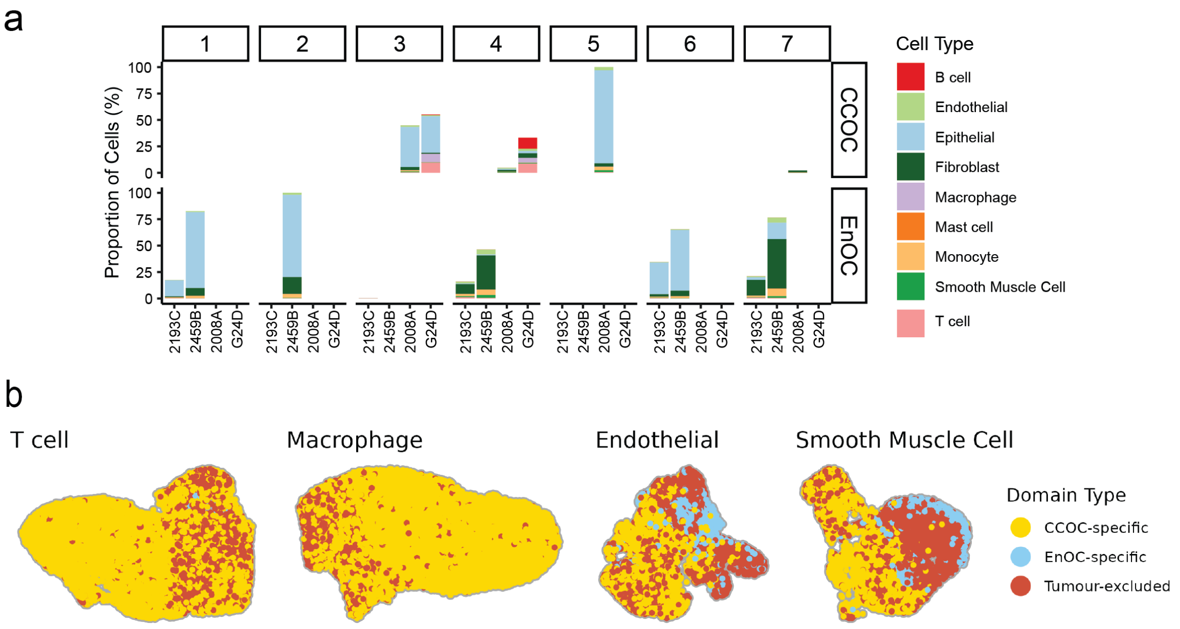


**Supplementary Figure 27.** (a) Proportion of cells from each cell type assigned to each of the seven integrative DOMINO domains across all samples. (b) UMAP embedding of stromal cell populations coloured according to assignment to CCOC associated, EnOC associated, or tumour excluded integrative DOMINO domains.

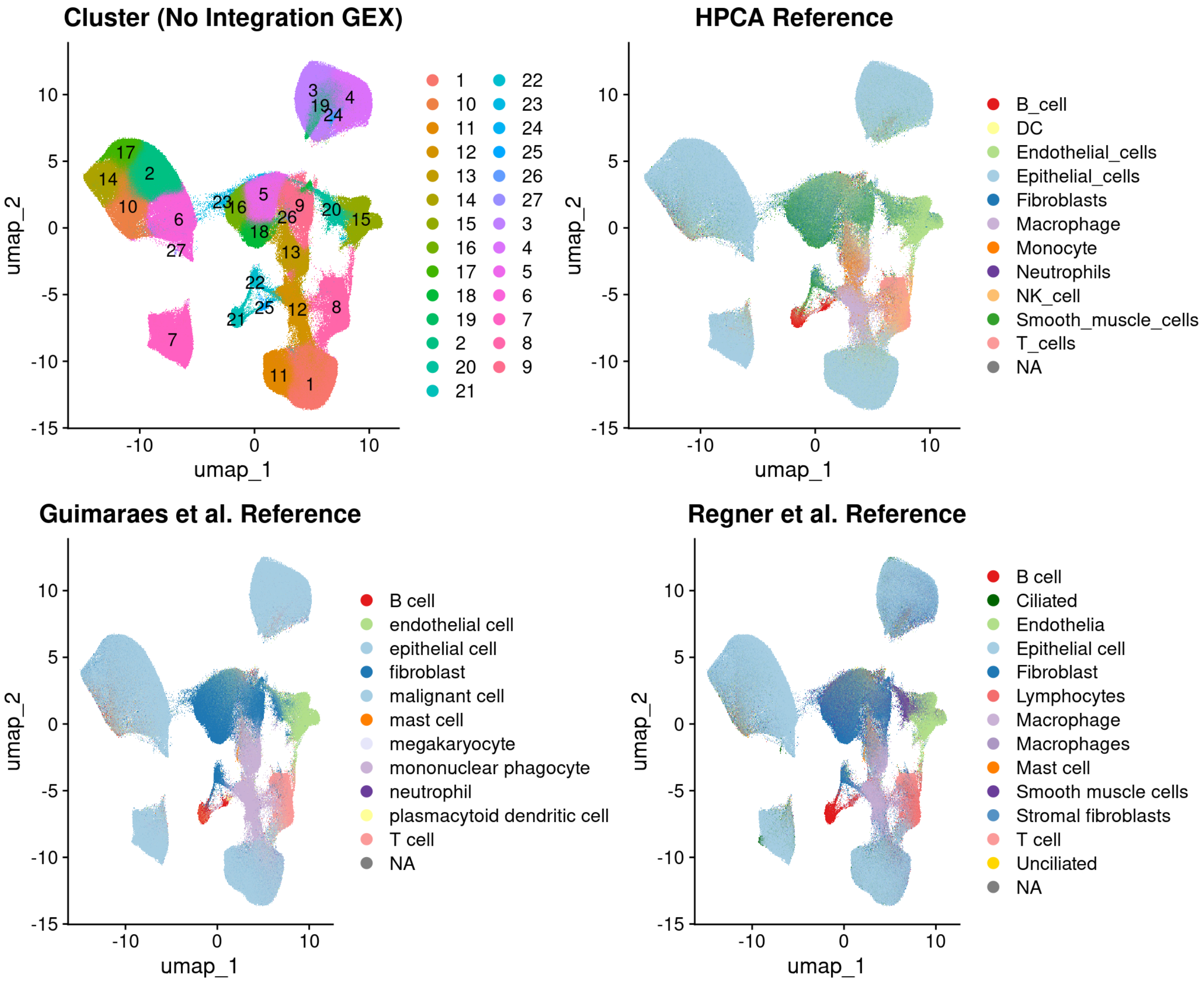


**Supplementary Figure 28.** UMAPs of all four samples generated from gene expression only. Cells are coloured by (i) gene expression-based clustering and (ii) SingleR cell type predictions using HPCA, Guimarães et al. and Regner et al. reference datasets.
